# Hope for Others: Research Results from the University of Pittsburgh Rapid Autopsy Program for Breast Cancer

**DOI:** 10.1101/2024.11.06.621982

**Authors:** Alexander Chih-Chieh Chang, Marija Balic, Tanner Bartholow, Rohit Bhargava, Daniel D. Brown, Lauren Brown, Adam Brufsky, Ye Cao, Neil Carleton, Amanda M. Clark, Morgan Cody, Kai Ding, Christopher Deible, Ashuvinee Elangovan, Julia Foldi, Daniel Geisler, Christine Hodgdon, Naomi Howard, Zheqi Li, Jie Bin Liu, Oscar Lopez-Nunez, Dixcy Jaba Sheeba John Mary, Olivia McGinn, Lori Miller, Kanako Mori, Geoffrey Pecar, Nolan Priedigkeit, Shannon Puhalla, Margaret Q. Rosenzweig, Partha Roy, Laura Savariau, Stephanie Walker, Hunter Waltermire, Abdalla M Wedn, Alan Wells, Megan E. Yates, Jennifer Xavier, Adrian V Lee, Steffi Oesterreich

## Abstract

Breast cancer affects 1/8 of women throughout their lifetimes, with over 90% of cancer deaths being caused by metastasis. However, metastasis poses unique challenges to research, as complex changes in the microenvironment in different metastatic sites and difficulty obtaining tissue for study hinder the ability to examine in depth the changes that occur during metastasis. Rapid autopsy programs thus fill a unique need in advancing metastasis research. Here, we describe our protocol and processes for establishing and improving the US-based Hope for OTHERS (Our Tissue Helping Enhance Research and Science) program for organ donation in metastatic breast cancer. As of August 2024, we consented 114 patients and performed 37 autopsies, from which we collected 551 unique metastatic frozen tumor samples, 1244 FFPE blocks, 90 longitudinal liquid biopsy samples and developed 14 patient-derived organoid and 8 patient-derived xenograft models. We report in-depth clinical and histopathological information and discuss extensive new research and novel findings in patient outcomes, metastatic phylogeny, and factors in successful living model development. Our results reveal key logistical and protocol improvements that are uniquely beneficial to certain programs based on identifiable features, such as working closely with patient advocates, methods to rescue RNA quality in cases where tissue quality may degrade due to time delays, as well as guidelines and future expansions of our program.

**Statement of Significance:** Rapid autopsy programs are unique research settings with huge potential for studying metastatic cancer, however, they have complex research challenges. Our work provides a valuable resource in advancing this field of research.

## Introduction

Breast cancer affects 1 in 8 women throughout their lifetimes^1^, with survival at five years averaging 31% for patients who have distant metastases^2^. Despite significant gains in breast cancer research and improvements in treatment in recent years, including the advent of CDK4/6 inhibitors and novel HER2-antibody drug conjugates, much work still remains to be done^3^. The most lethal mechanism of breast cancer is metastasis, which is responsible for the majority of cancer deaths, and addressing this challenge remains a critical focus of ongoing research efforts^4,5^.

However, metastasis is uniquely challenging to study, as it involves a complex interplay between genetic and epigenetic modifications related to immune and other environmental factors that are not easily captured in the laboratory setting. Clinical samples are urgently needed as mouse models and other laboratory techniques may not fully capture the complexity of human genetics and disease^6^. However, many clinical tissues that are biopsied are not routinely preserved for research purposes and provide a limited number of organs and sites that may not capture the full picture of metastasis^7,8^.

Autopsies thus provide crucial diversity in the tissues collected for research purposes, which leads to a more explicit understanding of the pathways taken during cancer metastasis^7^. Specific advantages include access to metastatic lesions that are challenging to biopsy, such as bone; access from normal tissues to study organ tropism as well as intra/inter-organ heterogeneity; larger amounts of tissue for in-depth molecular study/model development; and the collection of tissue after lines of therapy to study drug resistance – a major challenge in breast cancer treatment^9^.

The establishment of a tissue donation program at UPMC Magee Women’s Hospital was driven by patient requests within the Breast Cancer Program, a component of the NCI-designated UPMC Hillman Cancer Center and the Magee Women’s Hospital. Between 2008 and 2015, four patients with metastatic breast cancer nearing the end of their lives expressed a desire to contribute to scientific research through body donation^10^. While these initial requests were accommodated, the process lacked structure and organization. The development of this structured tissue donation program was motivated by the need to streamline the process, maximize the scientific value of donated tissues, and fulfill the wishes of patients who sought to contribute to the advancement of breast cancer research even after their passing.

The significant quantity and quality of tissue obtained from these autopsies, coupled with the recognition that serial tissue collection throughout metastatic breast cancer progression enhances the value of autopsy tissue, highlighted the need for a formalized and proactive tissue procurement program. This initiative aimed to gather tissue samples throughout the illness and to ensure efficient and timely tissue collection at death, thereby honoring the patients’ desire to leave a lasting impact on cancer research and, thus, on future patients suffering from the disease through improved understanding of breast cancer evolution, heterogeneity, and metastases.

At the program’s inception, no cancer-specific tissue autopsy procurement programs existed within our academic center. However, an existing rapid autopsy program for Idiopathic Pulmonary Fibrosis (IPF) patients served as a valuable model^10^. This IPF program helped identify crucial departments, personnel, and procedural workflows necessary for conducting autopsies effectively^11^, and other major logistical changes that need to be implemented. Hence, a major re-design was implemented in 2018, leading not only to exponential increases in consents and autopsies but also to increases in the quality and quantity of tissue collection and research progress.

This study presents a detailed analysis of the development, implementation, and outcomes of our rapid autopsy program, addressing logistical challenges and highlighting solutions. We report on diverse causes of death in metastatic breast cancer and emphasize the importance of systematic tissue collection, including the discovery of micro-metastases in grossly normal organs. Methodological advances in tissue preservation, particularly fixed sequencing technologies for RNA integrity in post-mortem samples, are discussed alongside the development of patient-derived organoids (PDOs) and xenografts (PDXs). Novel findings, such as the identification of an ESR1-ARNT2 fusion in metastatic samples from one patient, demonstrate the program’s potential to uncover new molecular features of metastatic breast cancer. Our experience provides valuable insights for improving rapid autopsy protocols and advancing metastatic cancer research globally.^12^

## Results

### Overall logistics of the HfO program from consenting to tissue processing

Our process for the Hope for OTHERS (Our Tissue Helping Enhance Research and Science) Tissue Donation Program illustrated in Figure 1 begins with the patient learning about and consenting to the program, which can occur years to days before their passing. Previous research from our group has shown that many patients are willing to discuss autopsy, but care providers must initiate such conversations^13^. As of August 2024, our average time from consent to death is 14-15 months, with a range of 0 to 53 months and a median of 10 months (Figure S1). Upon consent, we integrate the collection of longitudinal samples such as blood, ascites, and biopsies throughout the patient’s treatment journey flagging them in coordination with the clinical team and biobanking them where possible. These samples are logged and preserved via the Pitt Biospecimen Core, allowing for comprehensive temporal analysis of cancer progression.

**Figure 1:**
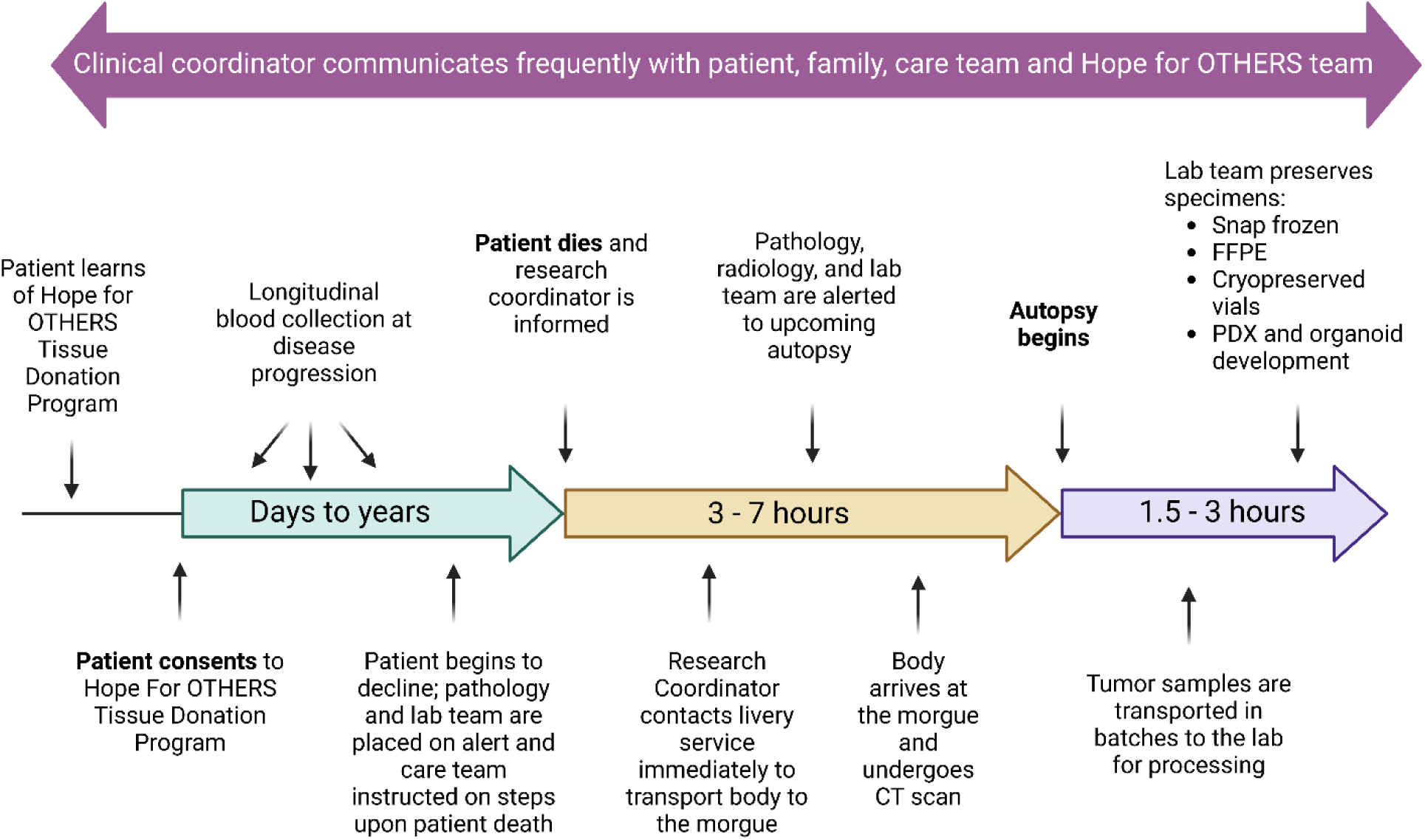
Study design and workflow of the HFO program. Diagram illustrating the study design and workflow of the HfO Tissue Donation Program.

As the patient’s condition declines, pathology and lab teams are alerted and prepared for potential tissue collection. Upon the patient’s death, the program enters a critical 3-7 hour window. The research coordinator is promptly informed and contacts a livery service to transport the body to the morgue. Simultaneously, pathology, radiology, and lab teams are notified about the impending autopsy. Once the body arrives at the morgue, we collect a post-mortem CT (computed tomography) scan.

The final phase, lasting 1.5-3 hours, involves the actual tissue collection. The autopsy begins, and the lab team works to preserve specimens using various methods. These include snap freezing, FFPE (Formalin-Fixed Paraffin-Embedded) preservation, cryopreservation, and the initiation of PDX (Patient-Derived Xenograft) and organoid development. Following collection, tumor samples are transported in batches to the lab for further processing.

As of August 2024, our program has consented 114 patients and completed 37 autopsies, averaging 5.5 autopsies per year since 2018. Our patients’ clinical characteristics (Table 1) reflect the general population demographics with ER+, PR+, HER2-being the most common molecular subtype. NST (no special type) is more prevalent than ILC (invasive lobular carcinoma) which also roughly approximates the frequency of ILC vs NST in the general population (∼15%) at 4/30 (∼13%) patients (Table 1). Stages at diagnosis ranged from 1A to 4. Our patients are most frequently diagnosed in stage 2A.

**Table 1:** Patient and Tumor characteristics of the HfO cohort as of August 2024.

| Clinical Characteristics |  |  |
| --- | --- | --- |
| Primary Molecular Subtype | ER+/HER2- | 22 (59.4%) |
|  | ER+/HER2+ | 6 (16.2%) |
|  | ER-/HER2+ | 1 (2.7%) |
|  | TNBC | 7 (18.9%) |
| Race | White | 35 (94.6%) |
|  | Black | 2 (5.4%) |
| Gender | Female | 36 (97.3%) |
|  | Male | 1 (2.7%) |
| Histological Subtype | NST | 33 (89.2%) |
|  | ILC | 4 (10.8%) |
| Stage at Time of Diagnosis | I - II | 19 (51.4%) |
|  | III | 9 (24.3%) |
|  | IV | 4 (10.8%) |
|  | Unknown | 5 (13.5%) |

Figure 2 presents a comprehensive timeline of treatments and outcomes for the 30 patients from the post-2018 autopsies [pre-2018 not included due to inconsistent data prior to the 2018 program re-design], each represented by a horizontal bar plotting their treatment and progression information, normalized by total duration from the patient’s initial diagnosis to the time of death. The first five squares color-code the characteristics of the patient for primary tumor molecular subtype, race, gender, histological subtype, and stage at the time of diagnosis.

**Figure 2:**
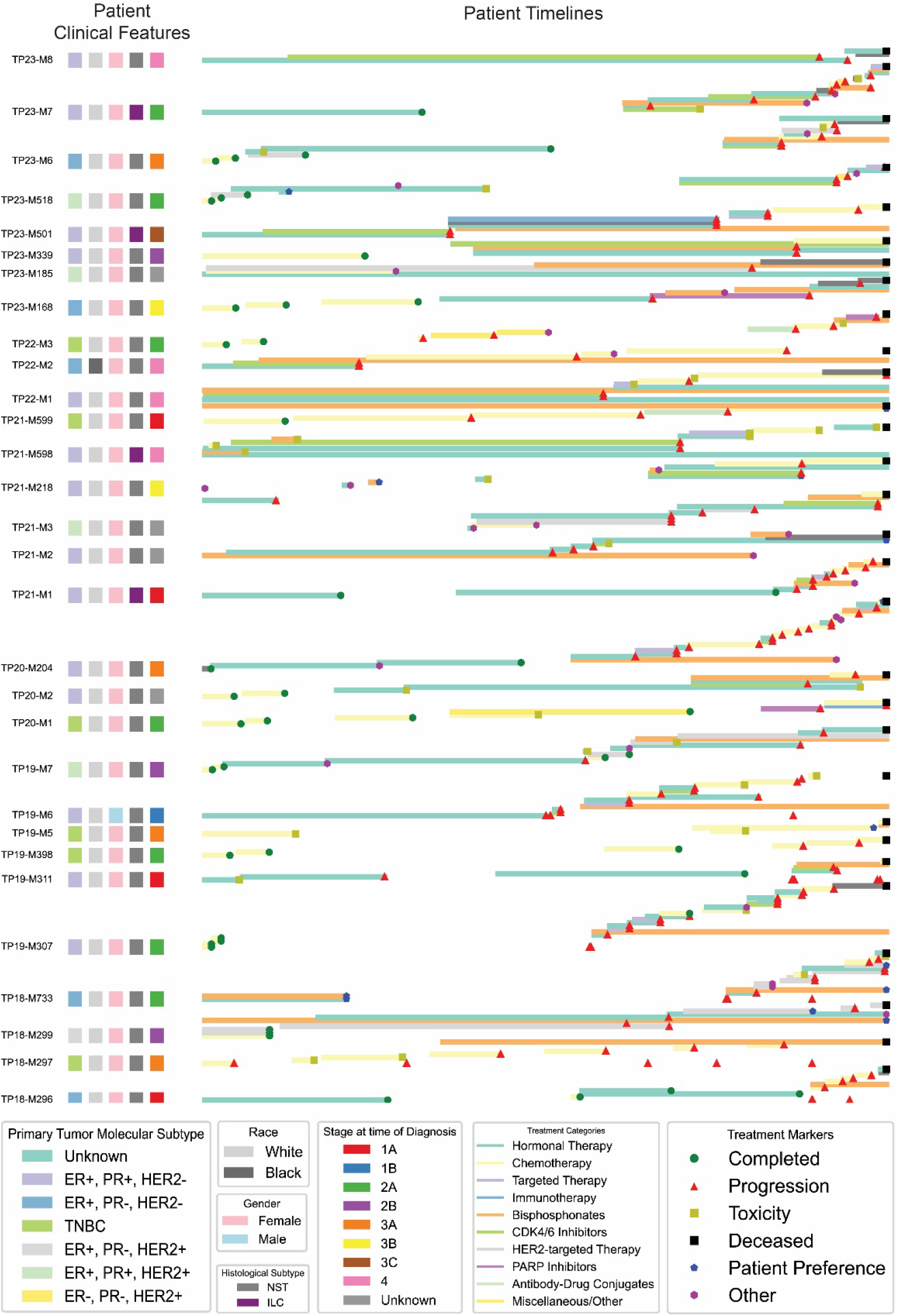
Summary of patients’ treatment timelines and clinical details. Mixture line and normalized timeline chart showing a summary of HfO program, including primary tumor molecular subtype, race, gender, histological subtype, stage at time of diagnosis (pathological if available, clinical if not), treatment markers and treatment categories up to August 2024.

Lastly, the treatment timeline uses color-coded bars to represent different therapies, including hormonal therapy, chemotherapy, targeted therapies, immunotherapy, and various inhibitors and conjugates. Simultaneous therapies are defined by vertically stacking bars. Our data shows that most of our patients, with some exceptions, follow similar treatment lines for their disease subtype. Symbols on the bars represent treatment markers, with an increasing density of red triangles later in treatment reflecting an increase in the progression rates in late-stage disease.

### Patient advocates play a crucial role in improving perceptions of the program

The development of rapid autopsy and organ donation programs for breast cancer research is a sensitive undertaking that requires careful consideration of ethical, emotional, and practical concerns. As we embarked on this initiative, it quickly became apparent that the perspectives of those most intimately affected by breast cancer—the patients themselves—were indispensable. Recognizing the delicate nature of discussions surrounding end-of-life care and post-mortem tissue donation, we realized that incorporating breast cancer advocates into our program was not just beneficial, but essential. These advocates, often breast cancer survivors and/or individuals with close ties to the breast cancer community, bring a unique and vital viewpoint to the table. Their involvement ensures that our approach remains patient-centered, addressing the concerns and honoring the wishes of those who might consider participating in such programs.

Hence, as an extension of our process, our group specifically incorporated a group of patient advocates with metastatic disease to represent patient voices on the leadership committee. This has led to several notable improvements, as discussed below.

This group coordinated a rebranding of our program to The Hope for OTHERS (Our Tissue Helping Enhance Research & Science; HfO) Tissue Donation Program. The new name was carefully chosen to reflect the altruistic nature of tissue donation and its critical role in advancing scientific understanding of breast cancer.

As part of this rebranding effort, we developed an independent, patient-focused website and created new materials such as brochures and pamphlets (Figure 3a). These resources were designed to provide clear, compassionate information about the program to potential participants and their families.

**Figure 3:**
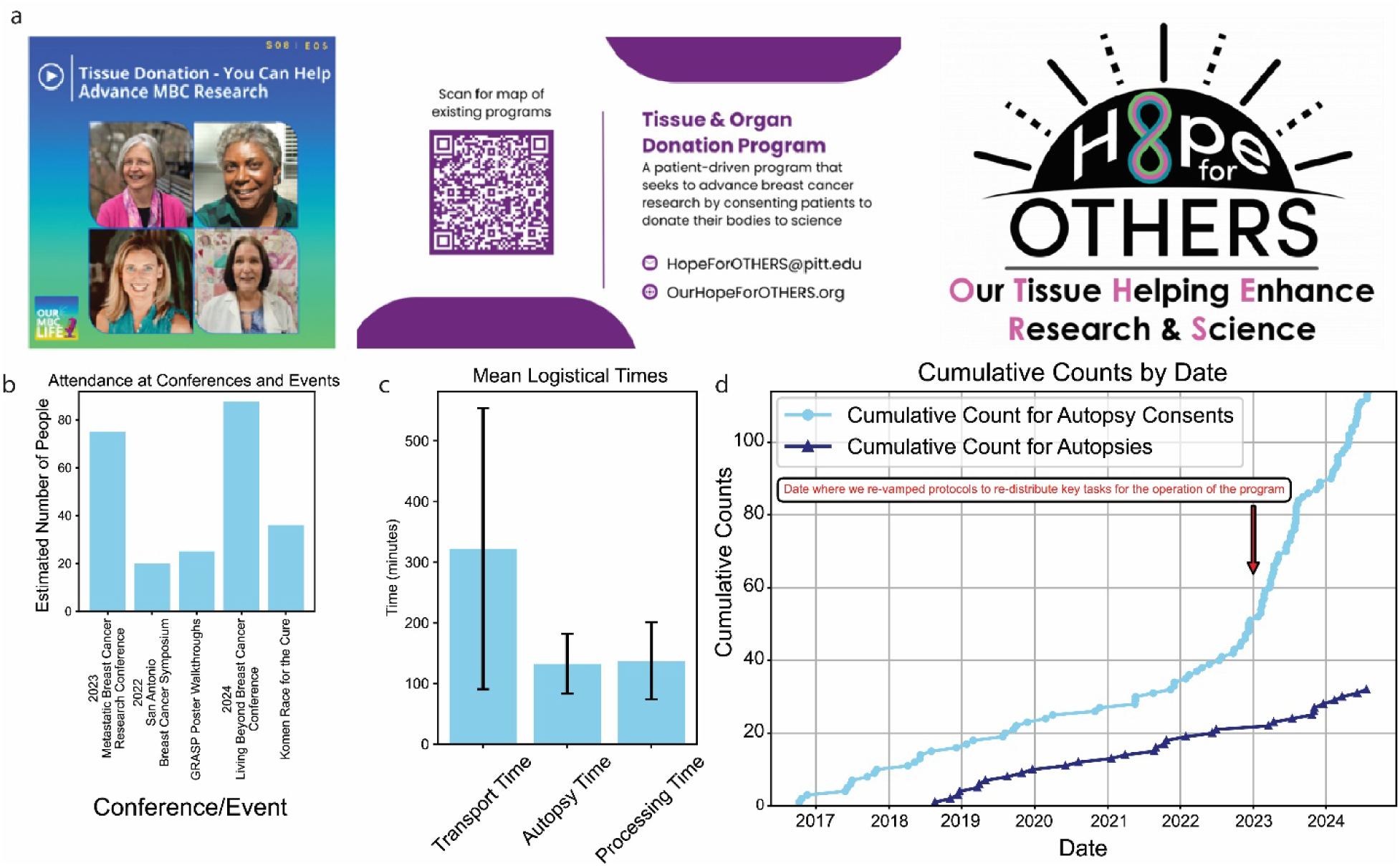
Examples of patient advocate media and figures demonstrating impact of key support staff. **a**. Promotion of tissue donation programs for advancing metastatic breast cancer (MBC) research via new media approaches such as podcasts, published media, and logo of the HfO program. Photos are of co-authors in this manuscript. **b**. Graph showing number of patients engaged at conferences/events. **c**. Bar plot shows the considerable standard deviation in mean transport time due to unique complexities within institutional and geographic contexts. **d**. Line graph shows the exponential increase in consent after an additional review of our operational protocols and rebranding (n = 114 consents, n = 34 autopsies [pre-2018 are not counted]). P-value from segmented regression 6.38E-31 for a change in consent rate slope post protocol review.

Our patient advocates have become integral members of the team, participating in our larger Hope for OTHERS meetings and holding additional meetings among themselves to discuss program improvements and outreach strategies. Their involvement ensures that patient perspectives are consistently represented in all aspects of the program. For example, we have recently participated in a podcast raising awareness of our program, which has been downloaded 330 times to date across 17 countries as of August 2024 (Figure 3a).

To increase awareness and engage with the broader breast cancer community, our program’s patient advocates have been actively presenting and distributing our materials (Figure 3a) at various regional and national conferences, such as the Metastatic Breast Cancer Research Conference 2024, Living Beyond Breast Cancer 2024, as well as advocacy events such as the 2024 Komen Pittsburgh More Than Pink Walk (Figure 3b).

Working closely with the advocate team and collaborating with groups across the country, we have seen a significant increase in interest in tissue donation for research purposes in metastatic breast cancer. Improving the perception of the program and its goals through frequent bidirectional interactions ensures that the patient perspective remains central to our efforts, furthering our mission of advancing breast cancer research through the normalization of patient donations.

### A clinical coordinator specifically dedicated to the HfO program assures communication and increases program efficacy

Rapid autopsy programs require engagement with numerous stakeholders, including but not limited to multiple clinical and basic departments, patient advocates, patients’ families, industry collaborators, researchers, regulatory offices, and funding agencies. The multi-layered complexities of the program require the commitment of and oversight by scientists who are truly vested in the success of the program, for example, those with a research focus on metastatic breast cancer, as is the case in our program.

An essential improvement of our program has been the addition of a dedicated clinical coordinator. The clinical coordinator is an integral part of the program, interacting with care providers, patients, and their families, pathologists on call, livery service, and the lab specimen processing team (Figure 1). While factors such as travel time for patient transport are outside of the coordinators control, we are only able to keep a narrow and consistent autopsy (Mean: 132.15 minutes, Standard Deviation: 49.23 minutes) and processing time (Mean: 137.03 minutes, Standard Deviation: 63.44 minutes) due to coordinator communication between the research lab and the autopsy team (Figure 3c), as well as maintain a steady increase in rates of consents due to their integrated and dedicated role in the clinic (Figure 3d).

We have established a structured meeting schedule to maintain program flexibility and continuous improvement. Biweekly meetings with the active operational group focus on case discussions, areas for improvement, and research progress monitoring. Bimonthly meetings involving all multidisciplinary team members and major stakeholders allow for sharing results and discussing larger-scale improvements, such as annual reviews of our standard operating protocols. A major re-design of roles and responsibilities, in 2023, by shifting more research responsibilities to dedicated research and autopsy coordinators, led to a dramatic increase in our rate of consents by the clinical coordinator due to a more focused and narrow scope as a result (P-value 6.38E-31) (Figure 3d).

### Comprehensive and Diverse Tissue Collection Enhances Longitudinal Metastatic Breast Cancer Research via Rapid Autopsy

At autopsy, we currently prioritize three collection modalities:

1. FFPE cassettes
2. Snap-frozen tissue for molecular analyses
3. Cryopreserved tissue for the development of patient-derived organoids (PDOs) and patient-derived xenografts (PDX)

As of August 2024, we have collected a sum total of 1244 FFPE blocks, with a median of 41 per patient (range of 7 – 69) (Figure 4a).

**Figure 4:**
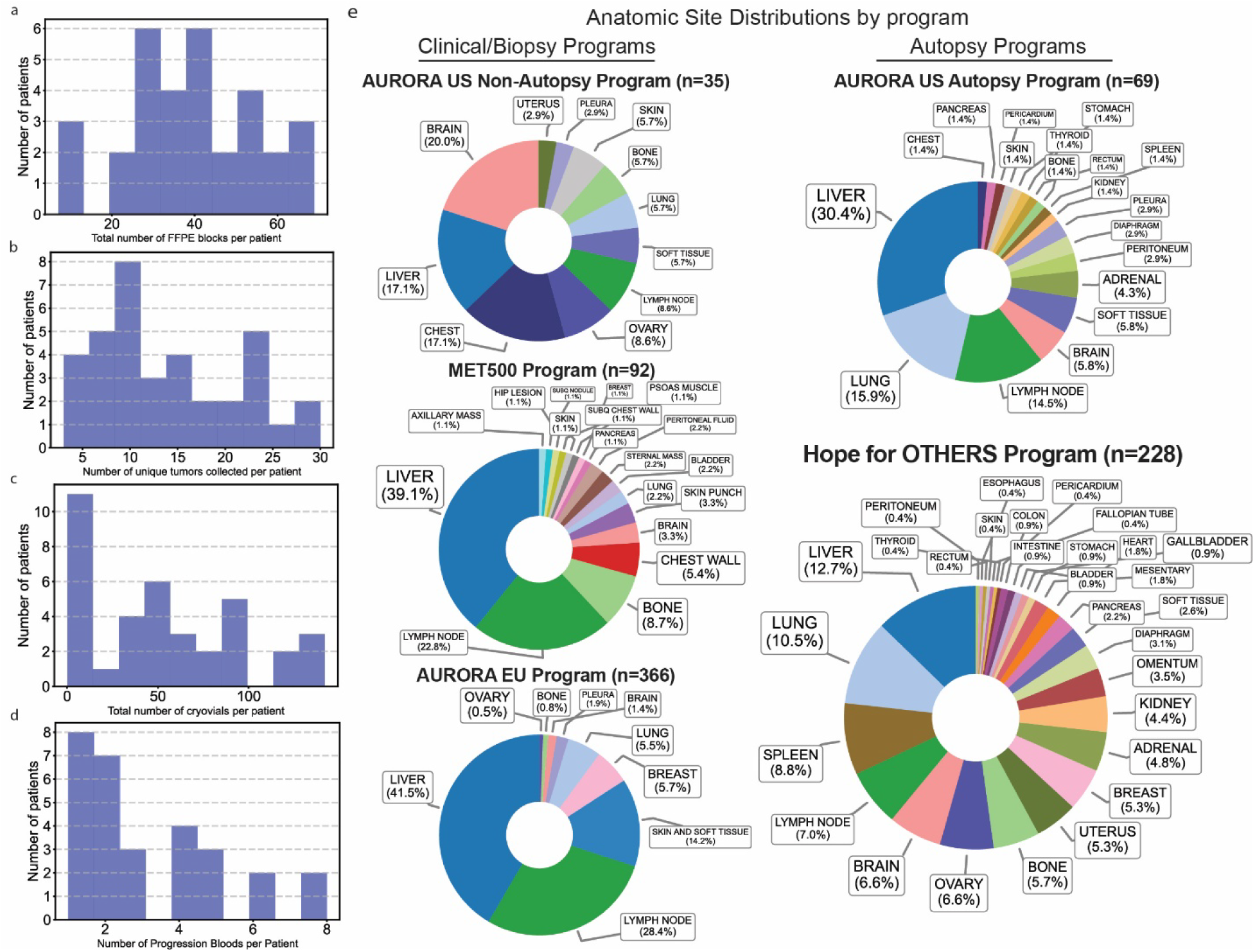
Summary of collected samples, and statistics on tissue diversity and counts in autopsy and non-autopsy settings. **a**. Histogram that summarizes statistics for FFPE. Total of 1244, median of 41, range of 7 to 69. **b**. Histogram that summarizes statistics for frozen tumors. Total of 511, median of 12, range of 3-27. **c**. Histogram that summarizes statistics for cryovials from autopsy. Total of 1952, median of 46 per patient, range from 0 to 142. **d**. Histogram that summarizes our longitudinal blood collections. **e**. Segment pie charts using data from the US Aurora, EU Aurora, MET500, and HfO reports showing the distinctly different range of tissues collected in autopsy and non-autopsy settings.

We have also collected 511 unique frozen tumors, with a median of 12 per patient (range of 3-27) (Figure 4b). A key strength of our program is the ability to access the original primary tumor samples for 25 of our cases (64%), despite sometimes being decades between primary surgery and death. This, combined with intermediate samples such as liver biopsies from the clinic, creates unique opportunities for longitudinal studies examining the evolution of metastatic breast cancer. Lastly, we have also collected 1952 cryovials, with a median of 46 per patient, and a range of 0 to 142 (Figure 4c).

Our research strategy also prioritizes the longitudinal capture of patient data, which can then be linked to future research studies; this includes both clinical data and, more importantly, liquid biopsy blood collections at each progression. These collections are a result of close coordination and collaboration with the medical oncology department and allow us to keep track of patient status, treatment progression, and other important clinical notes that might otherwise be missed and consolidate them for consistent formatting inclusive of the clinical context of each progression. For our 37 cases, we have collected 90 total, with a median of 2 per patient and a range of 1 to 8 progression blood collections (Figure 4d).

A primary improvement due to our autopsy process has been a noted increase in the diversity of tissue collected compared to previous studies in metastatic tissue. Even in studies that prospectively select for metastatic tissue, such as the 91 patients with metastatic breast cancer in the MET500 study, biopsies for liver and lymph nodes are overwhelmingly the majority of samples collected (61.9%)^14^ due to ease of access in the clinical setting (Figure 4e). Our results highlight the advantage of rapid autopsy programs to increase diversity of tissue samples collected.

As of August 2024, our program has collected from 228 organ sites, with fairly equal representation from many tissue sites, with liver (12.7%), lung (10.5%), and spleen (8.8%), making the top three showing relatively similar levels of collection. Furthermore, we also show increased diversity with rare sites of micro-metastases or local invasion such as thyroid, bladder, and diaphragm also being collected. We have collected from 29 total unique tissues compared to 11 (AURORA US Clinical Samples), 18 (MET500), and 11 (AURORA EU) from programs that collect using clinical biopsies (Figure 4e)^7,14,15^. This emphasis on comprehensive sampling via an autopsy approach has allowed us to capture a more complete picture of metastatic progression and organ involvement. All tissues are kept in −80 or −150 freezers and in duplicate both in our lab and at the Pitt Biospecimen Core to ensure backups in case of power outage or other system failures.

### Differential causes of death in breast cancer necessitate consistent and diverse tissue collection

Recent research has identified that causes of death in metastatic breast cancer are varied and require further examination^16^. To investigate this, we conducted a systematic review of our clinical records and autopsy reports after noting discrepancies between patient symptoms and gross clinically detected metastases. We focused specifically on lab values and medical notes from the last six months of life as well as the final autopsy report after death.

This analysis showed that the primary cause of death in the majority of patients enrolled in the HfO program was due to liver or respiratory dysfunction secondary to breast cancer metastasis (Figure 5a). However, while the majority (90%) of the liver pathologies were similar, with consistent signs of hyperbilirubinemia, portal hypertension, and cardiac strain, lung pathologies were much more diverse in the proximal cause, including but not limited to chronic kidney disease secondary to pulmonary congestion, disseminated intravascular coagulation, saddle pulmonary embolism, pneumonia, enhertu-related pneumonitis, and pleural effusions. Some, such as saddle pulmonary embolisms, have known or hypothesized causes currently under study^17^. Critically, many organ systems were revealed to have detrimental effects on others, such as the liver on the kidney due to breakdown of blood supply, or how pleural effusions or hepatojugular reflux could each cause cardiac strain. This illustrates how disentangling the specific cause of death in multisystem organ failure due to complications associated with metastatic breast cancer is a challenging task. Our data shows that causes of death in patients can vary in pathology and urgently demand further precise investigations into the underlying biology.

**Figure 5:**
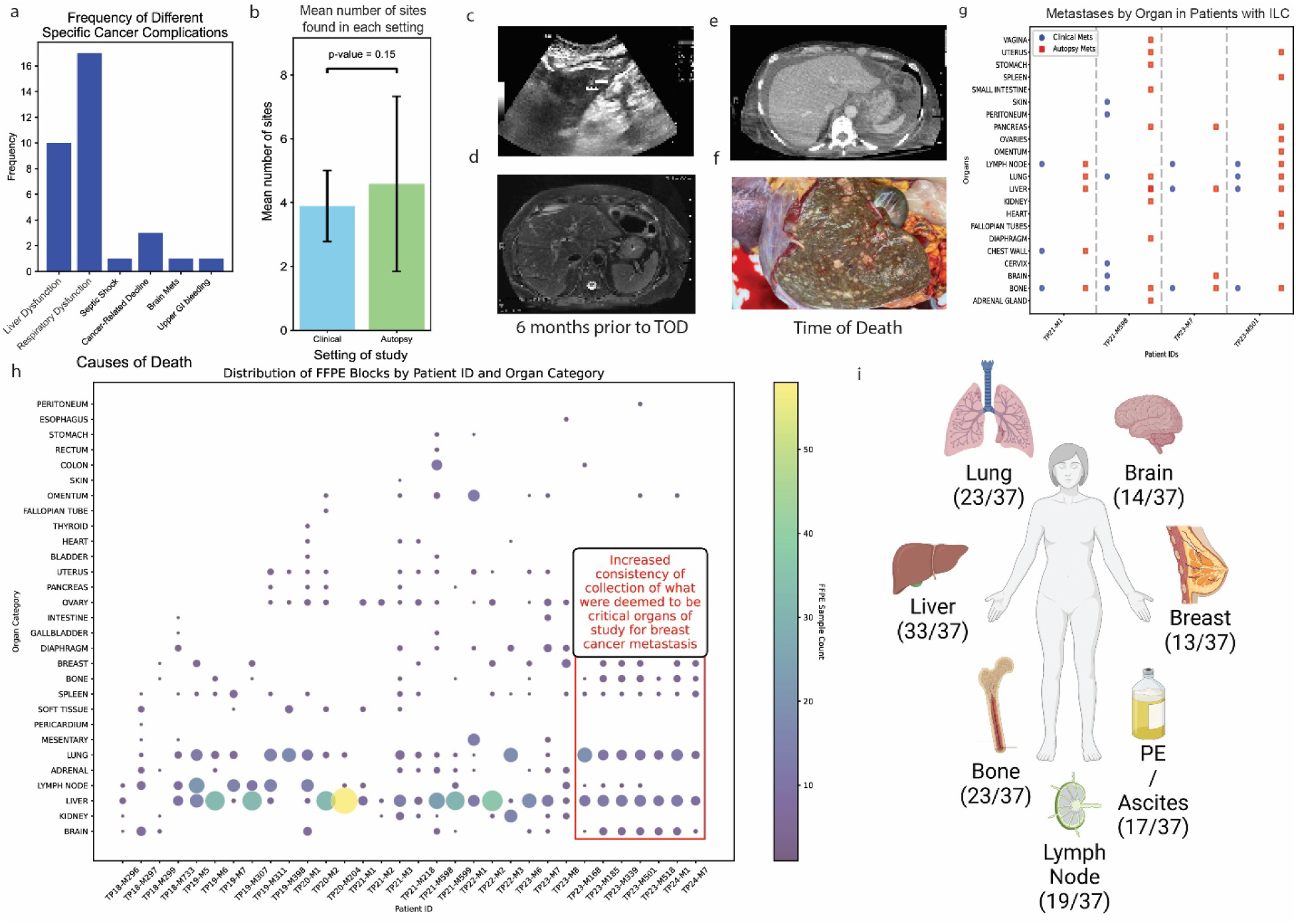
Diverse causes of death and subclinical metastases seen in autopsy settings, necessitating consistent collection. **a**. Bar plot showing the frequency of different causes of death in our program based on clinical note and autopsy report review (n = 37). **b**. Bar chart showing the mean increase in organs identified with metastases after careful pathological review on autopsy that were not identified in regular clinical monitoring, error bars are standard deviation. P-value 0.15 with paired t-test. Images of **c**. ultrasound and **d**. MRI in a patient with ILC showing the CT-undetectable liver metastases. **e**. CT image and **f**. autopsy image at time of death for the patient with ILC illustrating the discrepancy between a ‘normal’ CT and the organ status. **g**. Scatter plot showing the difference in metastases seen clinically and in autopsy for patients with ILC. Patients with ILC have much more spread in peritoneal tissues that are undetectable clinically. **h**. Scatter bubble plot showing our FFPE collection, red box highlights our improved protocol to increase consistency in grossly normal tissues. Color corresponds to size of bubble. **i**. Figure showing top 7 organ sites, collected whenever available, even if grossly normal under our revised protocol.

### Microscopic tumor metastasis is more common than expected and has different patterns between breast cancer histological subtypes

Often, the differences in metastatic organs can be microscopic. A review of all grossly normal organs from our patients showed that, on average, at least one additional organ from a patient may have micro-metastases not visible on clinical imaging (Figure 5b).

These have important implications for the journey of metastases and systemic responses to these metastases in different immune tissues. In some cases of ILC, patients were deemed to have ‘normal liver’ on CT repeatedly up until death, necessitating orthogonal approaches such as identification of lesions using ultrasound and MRI (magnetic resonance imaging); (Figure 5c, 5d) only to reveal at autopsy multiple metastatic lesions affecting the hepatic parenchyma despite consistently ‘normal’ CTs (Figure 5e, 5f) – a finding consistent with current ILC research and emphasizing the need for improved imaging as well as orthogonal methods of metastatic validation in addition to imaging^18^.

This kind of discrepancy was observed to be common in our patients with ILC, where numerous peritoneal organs often have metastases that are not visible clinically, as seen in Figure 5g. As a result, we made changes in our collection protocol (see Supplemental 1) that have resulted in increased standardized collection of lung, liver, brain, and bone in patients with NST and peritoneal organs and tissues in patients with ILC, with clear anatomical labels and records, increasing data representation overall for these diseases (Figure 5h). In addition, the need and desire to understand dormancy, especially in patients with late recurrences, such as in the case of ILC^19^, has prompted us to increase the collection of macroscopically normal tissue, such as bone, lung, and spleen, as well as sites considered to be ‘commonly’ involved in cancer metastasis – even if grossly normal (Figure 5i).

### Fixed single cell sequencing technologies are preferred for tissues with degraded RNA in the autopsy context

Previous research from Geukens et al. has shown that bulk RNA quality in tumor tissue decreases rapidly within hours of time of death^20^. In order to explore this and extend this work, we performed paired single nuclei sequencing from three liver samples using 3’ chemistry from frozen tissue that was snap frozen at autopsy, and using fixed flex technology from 10x Genomics on matched fixed tissues that were fixed and paraffin embedded at autopsy. Fixed flex uses multiple probes to identify fragments and multi-align probe signatures^21^.

Our results showed significant degradation of RNA from frozen tissues, which is in line with findings from previous research that RNA quality degrades in an autopsy setting^20^. At the same targeting of 8000 cells, on average, 7920 cells (SD: 1115) pass quality control for fixed sequencing, but only 6959 (SD: 585) pass quality control for frozen sequencing.

Additionally, fixed sequencing technologies can rescue some of the signals, resulting in an increased number of genes detected. Fixed sequencing detects an average of 2408 genes [95% CI 2387 - 2430] vs. 1454.77 [95% CI 1440.59 – 1468.95] in frozen sequencing. Fixed sequencing also resulted in increased total molecular counts with an average of 3158.05 [95% CI 3145.28 – 3170.81] total counts vs an average of 2357.83 [95% CI 2347.03 – 2368.63].

Lastly, we also see decreased mitochondrial contamination with an average of 0.65% mtDNA percentage [95% CI 0.64 – 0.65] vs 0.92% [95%CI 0.91 – 0.94] (Figure 6a).

**Figure 6:**
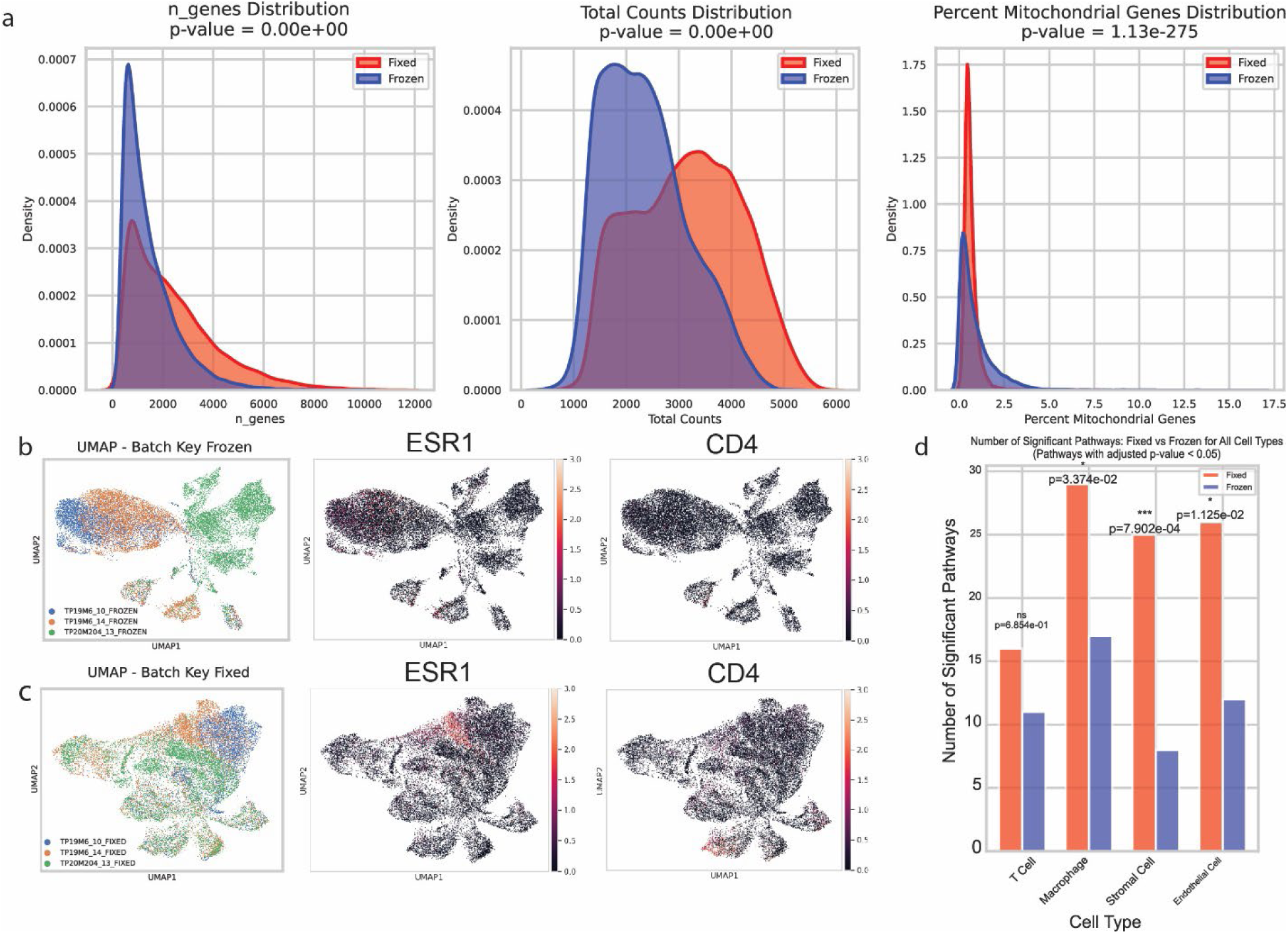
Single nuclei sequencing of paired frozen and fixed tissue samples. **a**. Side by side comparison of frozen vs fixed single nuclei sequencing data metrics shows significant improvement in fixed technologies with increase in genes detected, molecules detected, and less mitochondrial contamination. (n = 3) Run using 10x Genomics kit targeting 8000 cells. **b**. UMAP plot showing that frozen single nuclei sequencing lead to loss of signal causing failure of integration and removal of batch effects (n = 3), with diminished ESR1, and CD4 signal. **c**. Fixed single nuclei sequencing has better integration due to better signal recovery, with visible improvements in ESR1 and CD4 signal. PGR signal consistency across both sets shows that fixed technologies are not artificially introducing signal that isn’t there. **d**. Bar plot showing the number of Hallmark pathways that have adjusted p-value less than 0.05 after pathway analysis between fixed and frozen cells, showing that fixed tissue almost always has better pathway signal. P-value calculated using chi-square test.

Here, we see that fixed technologies also reduced noise, as these samples integrate better in the case of fixed technologies. They also show a wider variety of cellular populations with more significantly increased heterogeneity (Figure 6b and Figure 6c) – crucially Figure 6b reveals that even among samples processed together, degradation from the autopsy itself contributes to significant divergence in UMAP clustering and that in frozen technologies these differences are too large to be disentangled using Harmony, but are correctable in fixed sequencing in Figure 6c. We also see an increase in signal detection, as ESR1, and CD4 for example are higher detected (Figure 6c) when using fixed sequencing.

Downstream analysis is also impacted, in Figure 6d we integrate the samples and label by cell type. For quality assessment, we exclude the cancer cells specifically, as inherent cancer subclone heterogeneity and the expression of neuronal and stem-like markers common in cancer cells are confounding factors in this analysis comparing sequencing techniques, as single-cell sequencing analysis is unable to reliably deconvolute the number of subclones present and how much of the variation is due to inherent cancer heterogeneity vs. technique differences. We do not have such significant variations in the ground truth in other cell types which allows us to more accurately assess the quality of each sequencing technique. We then run side by side pathway analysis on cell type populations identified in both groups and show significant reduction in pathway activity detection in the Hallmark pathways.

While the same pathways are present in both fixed and frozen data in Figure S2, there is always a decreased gene set percentage in frozen data and a decreased pathway score in all four cell types, indicating worse signal quality (Figure S2).

### Time from death of patient to processing tissue is critical for developing living models that have the potential to reveals metastatic disease evolution

Through our HfO program, we have established additional corollaries for PDO and PDX generation. Specifically, we have been successful in generating 14 PDOs from 7 patients, and 8 PDX from 4 patients, covering a range of molecular and histological subtypes.

Across 27 attempts with 13 successes at developing PDOs, logistic regression revealed time to end of processing was a significant factor in organoid growth success, with the latest success in our tests being at 9 hours (Figure 7a) and a coefficient of −0.0132 for each additional minute of delay from time of death to end of processing (p = 0.027). We have also recently had significant success with 4 cryopreserved PDO developments out of 5 attempts and are currently accelerating new attempts from prior banked samples to unlock the potential of past collections further. To increase the possibility of success of organoid development, in 2023, our updated SOP (Supplemental 1) changed to submerge all organs whole in pre-chilled DMEM, drawing inspiration from organ transplant protocols^22^. Work to quantify the improvement as a result of this change is ongoing.

**Figure 7:**
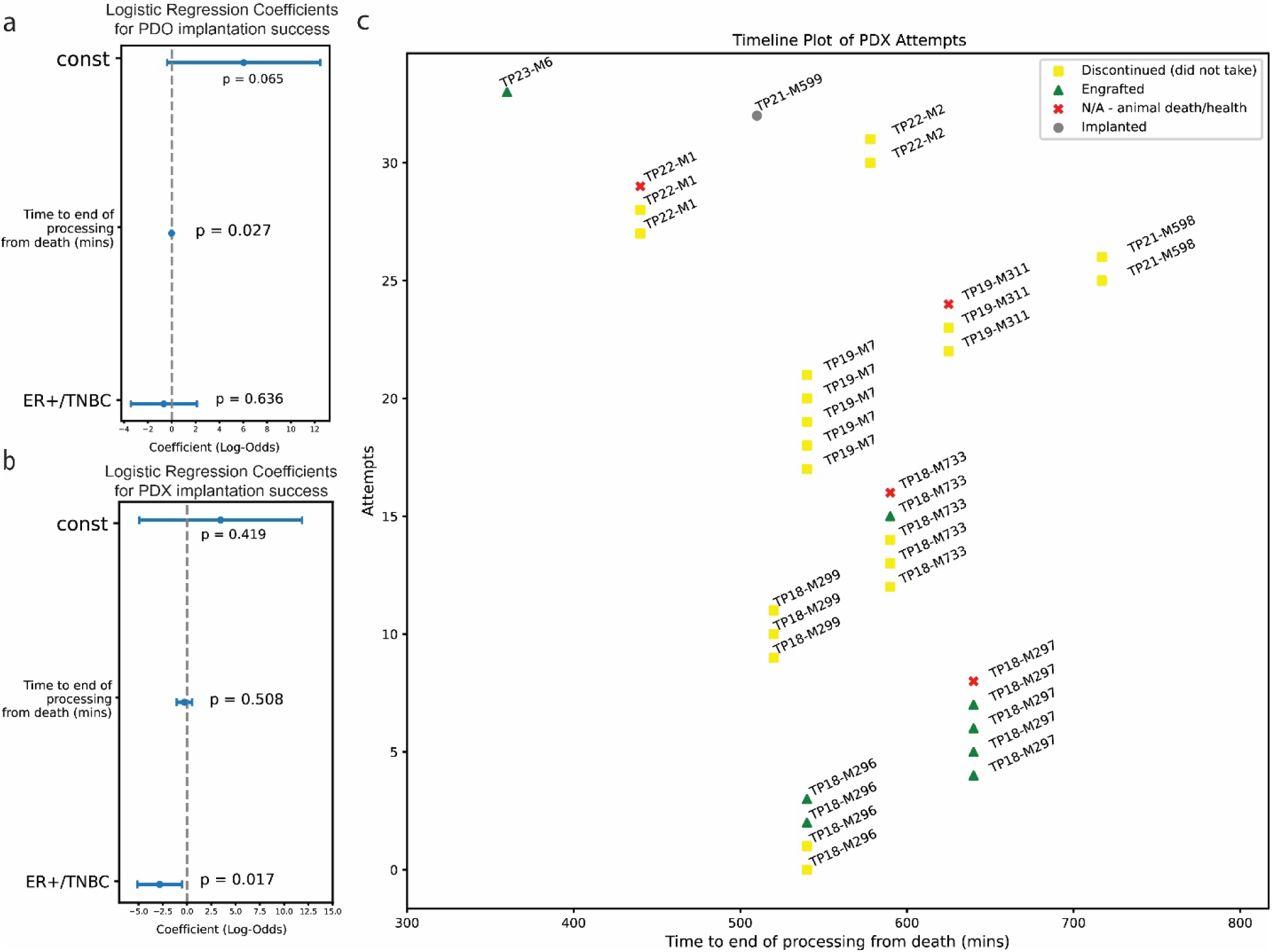
Logistic regression results for PDO and PDX generation success. **a**. Logistic regressions for PDO attempts (n = 27 attempts) show that time to end of processing is predictive of success for PDO success. **b**. Logistic regressions for PDX attempts (n = 34 attempts) show that TNBC lesions are more likely to be successful. **c**. There are also significantly different probabilities of success on an intra-patient basis, implying that tissue of origin or tumor cellularity may play a role (n = 34).

We are also collaborating with Champions Oncology on the generation of PDX models, with 11 patients, 34 attempts, and 8 successes, with no successes past the 10-hour mark after the time of death (Figure 7c). Lastly, logistic regression confirmed a commonly known factor in PDX success in triple negative vs. ER+ tumors, with coefficient of −2.8194 and p-value of 0.017, confirming prior findings that tumors that are ER+ have greater difficulty in engrafting^23^ (Figure 7b).

Furthermore, our collaboration with Champions has led to the generation of a custom protocol for excising tissue chunks for PDX implantation – noting that minced tissue is preferred for PDO generation, but chunked whole tissue is preferred for PDX generation (Figure S3).

Hence, careful evaluation of the number of each type of tissue preservation is essential to maximize downstream research options while operating with time constraints to transfer tissues in media and/or on ice as fast as possible.

### Novel research findings and the importance of collaboration with external partners

These PDX models have also given us unique opportunities for the study of metastatic progression. For example, we made the novel discovery of an *ESR1-ARNT2* fusion in a PDX model (Figure 8a) generated from a sample from a patient who participated in the HfO program. We then found the *ESR1-ARNT2* gene fusion to be ubiquitous among the metastatic tissues collected from our program for that patient (Figure 8b). We were subsequently able to cultivate a patient-derived xenograft organoid (PDXO) from this model (Figure 8c), and demonstrated that both the PDX and PDXO expressed the *ESR1-ARNT2* fusion gene (Figure 8d). We were able to express this fusion in cell-lines (Figure 8d), and are now studying it in greater detail. This then helps us build on previously published work from our lab as well that of other labs demonstrating that ESR1 fusions have unique activity and frequency in metastatic ER+ breast cancer tissue^24–26^.

**Figure 8:**
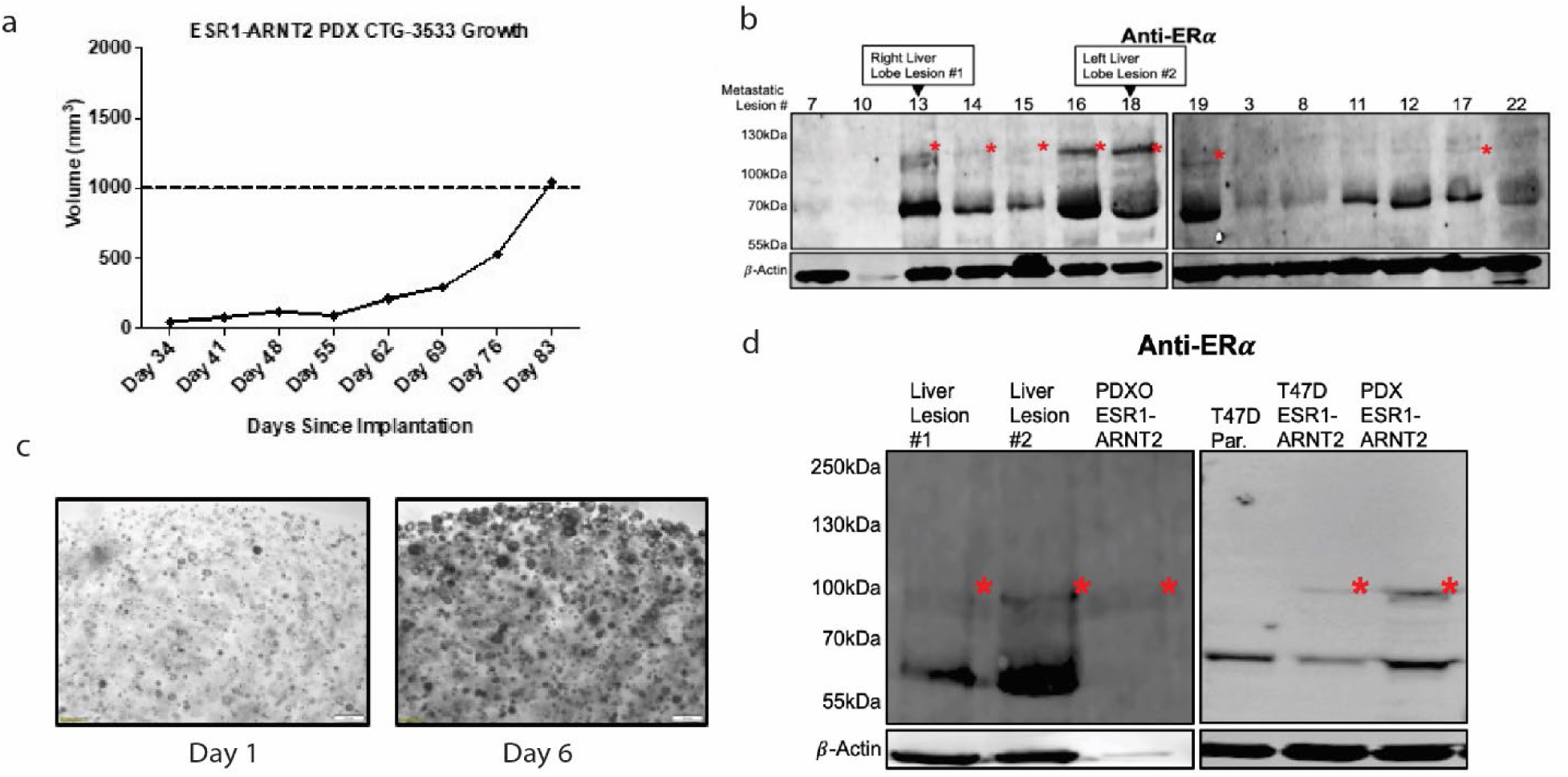
Validation of expression of the ESR1-ARNT2 fusion detected in HfO samples. **a**. Growth of PDX model CTG-3533. **b**. ER-immunoblot of TP18-M733 metastatic lesion protein samples. Red asterisks denote the expected size for the ESR1-ARNT2 fusion protein. *β*-actin serves as loading control in both blots, n=1. **c**, PDXO from CTG-3533 PDX can be cultivated from the PDX to extend back to in vitro experimentation. **d**. PDXO shows low levels of expression of the fusion construct and heterogenous signal. Sequencing also allows design of the fusion construct to be transfected into cells lines like T47D, as seen on ER-western blot. Red asterisks denote the expected size for the ESR1-ARNT2 fusion protein. *β*-actin serves as loading control in both blots, n=1.

The acceleration of our consent and autopsy progress has also been reflected in our research efforts. As of August 2024, we now have multiple projects in progress, including but not limited to research efforts looking at dormancy, genomic structural evolution, leptomeningeal metastasis, expansion of PDX model development to look at specific drug resistance models, ESR1 mutant effects in the liver microenvironment, and international collaborations at the clinical and molecular history of changes in breast cancer subtypes such as ILC with the UPTIDER program^20^. We expect critical discoveries in the coming years from the growing number of autopsy programs including ours. It’s important to note that the nature of these programs is highly dependent on the rate of sample collection and quality of data collected for research productivity and thus requires a certain degree of establishment before research output can catch up, which we have only recently managed to reach. This kind of investment requirement has significant funding and research planning implications.

## Discussion

The Hope for OTHERS program represents a breakthrough in metastatic breast cancer research. Our findings highlight the importance of rapid autopsy programs for understanding metastatic progression, while revealing challenges requiring multidisciplinary collaboration. Many programs fail without securing necessary support from the start, explaining why initiatives like ours remain rare globally^12^.

### Programmatic Improvements and Their Impact

A dedicated clinical coordinator has been crucial to our success by facilitating stakeholder communication and streamlining processes, resulting in increased patient consent and completed autopsies. The participation growth (Figure 3d) demonstrates this approach’s effectiveness as a potential model for other institutions.

Collaboration with patient advocates transformed our program through improved branding, better communication materials, and increased public awareness. This patient-centric approach has boosted participation while ensuring our research aligns with patient needs.

### Challenges and Solutions in Program Logistics

Transportation time challenges (Figure 3c) emphasize the need for flexible protocols in rapid autopsy programs. Our experience shows population density significantly impacts efficiency. Institutions with similar geographical challenges may benefit from adapting their protocols accordingly.

Our interdepartmental collaborations, particularly with Pathology and Radiology, have been essential in maintaining short time-to-autopsy windows and enhancing data collection quality, demonstrating the importance of an institution-wide approach.

### Scientific Insights and Methodological Advances

Our findings regarding the diverse causes of death in metastatic breast cancer (Figure 5a) underscore the complexity of the disease and the need for comprehensive tissue collection protocols. The discovery of micro-metastases in grossly normal organs (Figure 5b) highlights the importance of systematic sampling, even in apparently unaffected tissues. This approach has particular relevance for specific subtypes like ILC, where peritoneal (and other) metastases may be clinically occult (Figure 5g, 5h) and dormancy is a very critical yet unresolved issue.

Other improvements include rapid cooling of organs in medium to slow metabolic activity and molecular degradation^20^; consistent collection of all major organs even if grossly normal (Figure 5h); prioritization of tissues of key research interest, such as the leptomeninges.

The development of subtype-specific collection protocols (Figure 5h) represents a significant methodological advance. This tailored approach ensures more consistent and relevant tissue collection, potentially leading to more robust and representative datasets for future studies.

Our exploration of RNA quality preservation techniques (Figure 6a-d) provides valuable insights for researchers facing similar challenges with post-mortem tissue quality. The superior performance of fixed sequencing technologies in preserving RNA integrity and cellular heterogeneity information could inform future methodological choices in single-cell studies using autopsy tissues. Fixed sequencing technologies clearly decrease signal loss, as signal integration and removal of batch effects are much clearer in fixed analysis. We suspect that part of the batch effect is from the continued RNA degradation during extraction and lysis of cells to isolate single nuclei, while fixed RNA is more stable, and any FFPE artifacts are rescued by the redundancies in probe signals. In situations where tissue quality may be subpar or RNA quality is expected to be degraded due to autopsy-related factors, fixed sequencing technologies are preferred in single-cell applications. Fixed sequencing technologies, however, are not perfect, as we do note that certain signals, such as CD8, are not detectable in either set of analyses, and there is still significant RNA degradation compared to fresh single-cell sequencing collected in optimal conditions such as surgical resections^27^.

The time-sensitive nature of PDO model generation from autopsy tissues (Figure 7a) offers crucial guidance for researchers aiming to develop living models from rapid autopsy programs. These findings can help optimize tissue processing protocols and set realistic expectations for model development success rates. Important to note is the high variability in PDX success even within the same patient, implying significant factors other than time in PDX development and ER+ status that need to be further optimized. Work to establish the effect of tissue type is currently ongoing. The range of successes within patients also indicates that tissue quality based on blood supply, technique and other similar factors could significantly impact the success of PDX generation (Figure 7c).

### Novel Findings and Future Directions

The identification of the ESR1-ARNT2 fusion (Figure 8b) exemplifies the potential of rapid autopsy programs to uncover novel molecular features of metastatic breast cancer. This finding, along with our ongoing studies on ESR1 fusion functions, demonstrates how rapid autopsy programs can drive forward our understanding of treatment resistance and metastatic progression. Both models are valuable and available to collaborators for further research, emphasizing the importance of models developed from programs like ours.

The diverse range of ongoing projects stemming from our program, including studies on dormancy, leptomeningeal metastasis, and rare subtypes like ILC, showcases the broad impact of comprehensive rapid autopsy programs on breast cancer research and the potential for new discoveries previously unknown.

### Limitations and Future Considerations

Despite our successes, we acknowledge several limitations. The single-institution nature of our study may limit the generalizability of some findings. Specifically, inter-institutional variations in protocol may also lead to differences in downstream research results^12^. Additionally, while we’ve made strides in reducing time-to-autopsy, further improvements could enhance tissue quality and model generation success rates.

Future directions for our program include expanding collaborations with other institutions to increase sample diversity and validate our findings across different populations. We also aim to refine our tissue collection and processing protocols further based on emerging technologies and research priorities.

In conclusion, the Hope for OTHERS Tissue Donation Program demonstrates the profound impact that well-designed rapid autopsy programs can have on advancing metastatic breast cancer research. By sharing our experiences, challenges, and solutions, we hope to contribute to the broader effort of improving rapid autopsy protocols, increase patient enrollment and ultimately advancing our understanding of metastatic breast cancer biology.

## Methods

All dates and statistics are as of a freeze date of August 1^st^, 2024.

### Operating Protocol

Please see attached Supplemental 1.

### Frozen single nuclei extraction

#### Reagents and Buffers

Nuclei isolation was performed using the following reagents: Trizma® Hydrochloride Solution (1M, pH 7.4; Sigma T2194), Sodium Chloride Solution (5M; Sigma 59222C), Magnesium Chloride Solution (1M; Sigma M1028), Nonidet™ P40 Substitute (Sigma 74385), Phosphate-Buffered Saline (PBS) with 10% Bovine Albumin (Sigma SRE0036), and Protector RNase Inhibitor (Sigma 3335399001).

Two buffers were prepared:

Lysis Buffer (TST): Composed of 1X ST buffer (10 mM Tris-HCl, 146 mM NaCl, 21 mM MgCl2, 1 mM CaCl2), 0.03% Tween 20, and 0.01% BSA.

Wash Buffer: Prepared with 1% BSA, 0.2 U/μL RNase Inhibitor in PBS.

All buffers were pre-chilled on ice or at 4°C before use.

Frozen tissue samples were minced on dry ice and transferred to 1.5 mL microcentrifuge tubes, with a sample volume not exceeding 500 μL.

500 μL of chilled Lysis Buffer was added to each sample. Tissues were homogenized on ice using a Dounce homogenizer (Fisher 12-141-363) with 10-20 strokes over a 5-minute period.

An additional 500 μL of Lysis Buffer was added, and samples were incubated on ice for the remainder of the 5-minute period, with intermittent mixing.

Homogenates were filtered through a 70 μm-strainer mesh, and the flow-through was collected in a polystyrene round-bottom FACS tube.

The filtrate was transferred to a 1.5 mL tube, and 500 μL of Wash Buffer was added. Samples were centrifuged at 500 g for 5 minutes at 4°C.

The supernatant was carefully removed, and the pellet was resuspended in 500 μL of Wash Buffer. This washing step was repeated once.

After the final wash, nuclei were resuspended in Wash Buffer and counted. The suspension was adjusted to a final concentration of 1000 nuclei/μL.

If necessary, an additional filtration step using a 40 μm Flowmi filter was performed to remove any remaining debris.

Samples were kept on ice and then sent to the Single Cell Sequencing Core at the University of Pittsburgh, targeting 8,000 cells for downstream sequencing.

### Fixed single nuclei extraction

For FFPE single nuclei extraction, we used a modified version of the snPATHO-seq protocol provided to us by the lab of Luciano Martelotto^28^.

#### Reagents and Equipment

Reagents included Ethanol (Decon Laboratories #2701), Xylene (Epredia, 6601), Nuclease-Free water (Invitraogen, AM9938), 1x Phosphate Buffer Saline (Ca2+ and Mg2+ free) (Corning, 21-031-CV), Liberase TM (Millipore Sigma, 5401119001), RPMI1640 (Gibco), 10% BSA (),TST Buffer and Wash buffer (See above in Frozen Single Nuclei extraction). Equipment used included a Thermomixer with adjustable shaking (Eppendorf) and a refrigerated centrifuge.

Nuclei Isolation Procedure

2-4 tissue sections (25 μm-thick) or punches were collected and stored at 4°C if not used immediately.

Paraffin was removed by washing sections three times with 1 mL Xylene for 10 minutes each. Samples were rehydrated through an ethanol gradient (100%, 70%, 50%, 30%) for 1 minute each.

Samples were washed once with 1× PBS + 0.5 mM CaCl2.

Tissue digestion was performed in 1 mL RPMI1640 supplemented with Liberase TM (1 mg/mL), for 60 minutes at 37°C with shaking at 800 rpm.

After digestion, 400 μL of TST Buffer was added, mixed, and centrifuged at 850 x g for 5 minutes at 4°C.

The pellet was resuspended in 250 μL TST buffer containing 2% BSA and 1 U/μL RNAse Inhibitor, then homogenized using a Dounce homogenizer (10-20 strokes).

An additional 750 μL of the EZ Lysis buffer mixture was added, followed by further disaggregation by pipetting up and down and incubation on ice for 5 minutes.

The sample was filtered through a 70 μm PluriStrainer and centrifuged at 850 x g for 5 minutes at 4°C.

Nuclei were washed twice with wash buffer and resuspended in PBS 0.5x + 0.02% BSA, put on ice and delivered to the Single Cell Sequencing Core or stored in cryopreservation (see below).

Samples were then processed for Chromium X run using Chromium Fix RNA Profiling (10x Genomics) following the manufacturer’s protocol.

For cryopreservation, samples were supplemented with Enhancer solution (10x Genomics) and 0,22um filtered 10% Glycerol provided by the Single Cell Sequencing Core, incubated on ice for 10 minutes, and stored at −80°C.

This method was optimized for the preparation of nuclei suspensions from formalin-fixed, paraffin-embedded (FFPE) tissue samples for single-nucleus RNA sequencing applications.

### Patient Derived Xenograft Development

Tumor chunks were excised from the organ, with a slice down the middle for increased media perfusion. Samples were then shipped same day to Champion Oncology lab for implantation. For cryopreserved tissue, chunks 0.5 cm cubed in size were excised and frozen in freezing media (See Supplemental 1 for details). These were then shipped frozen to Champions Oncology overnight when required.

### Organoid generation and culture

Patient-derived organoids (PDOs) were generated from consented primary human breast cancer tissue from the Pitt Biospecimen Core in accordance with Institutional Review Board protocol STUDY22030183 by the Institute for Precision Medicine according to established protocol (Sachs et al 2018), with the addition of β-estradiol to the medium. Briefly, tumors were digested with collagenase (Sigma C9407) on a rotator, sheared, filtered, and embedded in Cultrex RGF Basement Membrane Extract (R&D Systems™ 353301002) in 24-well non-treated plates (Fisher 12-566-82). Media replaced every 2-3 days and PDOs passaged every 2-4 weeks.

### Organoid Growth Assay

PDOs were dissociated using 0.25% trypsin, washed in Advanced DMEM/F12, seeded at 20,000 cells per well in 96-well round bottom plates (Corning 353227), and cultured in standard growth media (250 ng/ml Recombinant Human R-Spondin-3, 5 nM Recombinant Human Heregulin β-1, 5 ng/ml Recombinant Human KGF (FGF-7), 20 ng/ml Recombinant Human FGF10, 5 ng/ml Recombinant Human EGF, 100 ng/ml Recombinant Human Noggin, 500 nM A 83-01, 5 mM Y-27632, 500 nM SB 202190, 1X B-27 Supplement, 1.25 mM N-Acetyl-L-cysteine, 5 mM Nicotinamide, 50 mg/ml Primocin, 10 mM HEPES, 1X GlutaMAX, 100 U/ml Antibiotic-Antimycotic, and 1X Advanced DMEM).

### Computational methods

Frozen single nuclei sequencing data were aligned using CellRanger 7.1.0 while Fixed Flex single nuclei sequencing was aligned using Multiranger 7.1.0. Standard Seurat Recipe Preprocessing was used and samples were then Harmony integrated for comparison. Celltype assignments were done using GSEAPY CellMarker2024 followed by manual review.

### Data Access Statement

All single-cell sequencing data will be made available on NCBI GEO at time of publication.

### Western Blot Methods

Cellular protein lysates were harvested utilizing RIPA buffer (50mM Tris pH 7.4, 150mM NaCl, 1mM EDTA (Thermo Fisher Scientific #15-575-020), 0.5% Nonidet P-40 (Sigma Aldrich #74385), 0.5% sodium deoxycholate, 0.1% SDS) supplemented with 1X HALT protease and phosphatase cocktail (Thermo Fisher #78442). Samples were vortexed, probe sonicated for 15 seconds (20% amplitude, Ultrasonic Processor GEX130) and centrifuged at 14,000rpm at 4 °C for 15 minutes. Protein concentration was assessed using the Pierce Bicinchoninic acid (BCA) protein assay (Thermo Fisher #23225). Unless otherwise stated, 50μg of each protein sample was run on a 10% SDS-PAGE gel followed with a 90V transfer at 4 °C for 90 minutes to a PVDF membrane (Millipore #IPFL00010). Membranes were blocked for one hour with Intercept PBS blocking buffer (LiCor #927-40000) at room temperature with rocking. Antibody probing was performed overnight at 4 °C with rocking: ER*α*, clone 60C (Millipore #04-820, RRID:AB_1587018); HA (C29F4) (Cell Signaling Technologies #3724, RRID:AB_1549585); *β*-actin (Millipore Sigma #A5441, RRID:AB_ 476744). After removal of primary antibodies, blots were wash with 1X PBSTween 20 (0.1%) for 15 minutes, three times. Secondary antibodies were applied for a one-hour room temperature incubation (1:10,000; anti-mouse 680LT (LiCor #925-68020); anti-rabbit 800CW (LiCor #925-32211)). Imaging of membranes was performed on the LiCor Odyssey CLx Imaging system.

## Supporting information

Supplemental 1 - Copy of our SOP

## Acknowledgements

We would like to thank the patients and their family and friends who believe in the program, and have contributed to it in many ways. We are forever grateful to those patients who consented to participate in the program.

We would like to thank the UPMC Cancer Registry, especially Vonda Mazzarella. Special thanks goes to many members of the Department of Pathology who have contributed to the program.

The authors would like to thank the Institute for Precision Medicine (IPM), a partnership of the University of Pittsburgh and UPMC, for providing the patient-derived breast cancer organoids used in these studies. Single Cell Gene Expression Flex was performed in the University of Pittsburgh Single Cell Core Facility (RRID:SCR_025110) and services and instruments used in this project were graciously supported, in part, by the University of Pittsburgh, the Department of Medicine.

We would like to acknowledge Champions Oncology as a partner for the development of PDX models for our program and look forward to continued advancements in model development.

We would also like to thank other peer organ donation and rapid autopsy programs for their collaboration and advice, and we look forward to a continued expansion in the global community of such programs to advance cancer research.

## Ethics Statement

All research involving human participants was conducted with informed consent obtained according to the ethical guidelines of the University of Pittsburgh Institutional Review Board under STUDY19060376.

## Funding Statement

The HfO program has been supported by the Magee Womens Research Institute and Foundation, and Susan G Komen Foundation. This project used the UPMC Hillman Cancer Center and Tissue and Research Pathology/Pitt Biospecimen Core shared resource which is supported in part by award P30CA047904.

Work performed in the Pitt Biospecimen Core (RRID:SCR_025229) and services and instruments used in this project were supported, in part, by the University of Pittsburgh, the Office of the Senior Vice Chancellor for Health Sciences.

This research was supported in part by the University of Pittsburgh Center for Research Computing through the resources provided. Specifically, this work used the HTC cluster, which is supported by NIH award number S10OD028483.

## Supplemental 1

A copy of our SOP for autopsy collection.

**Figure S1:**
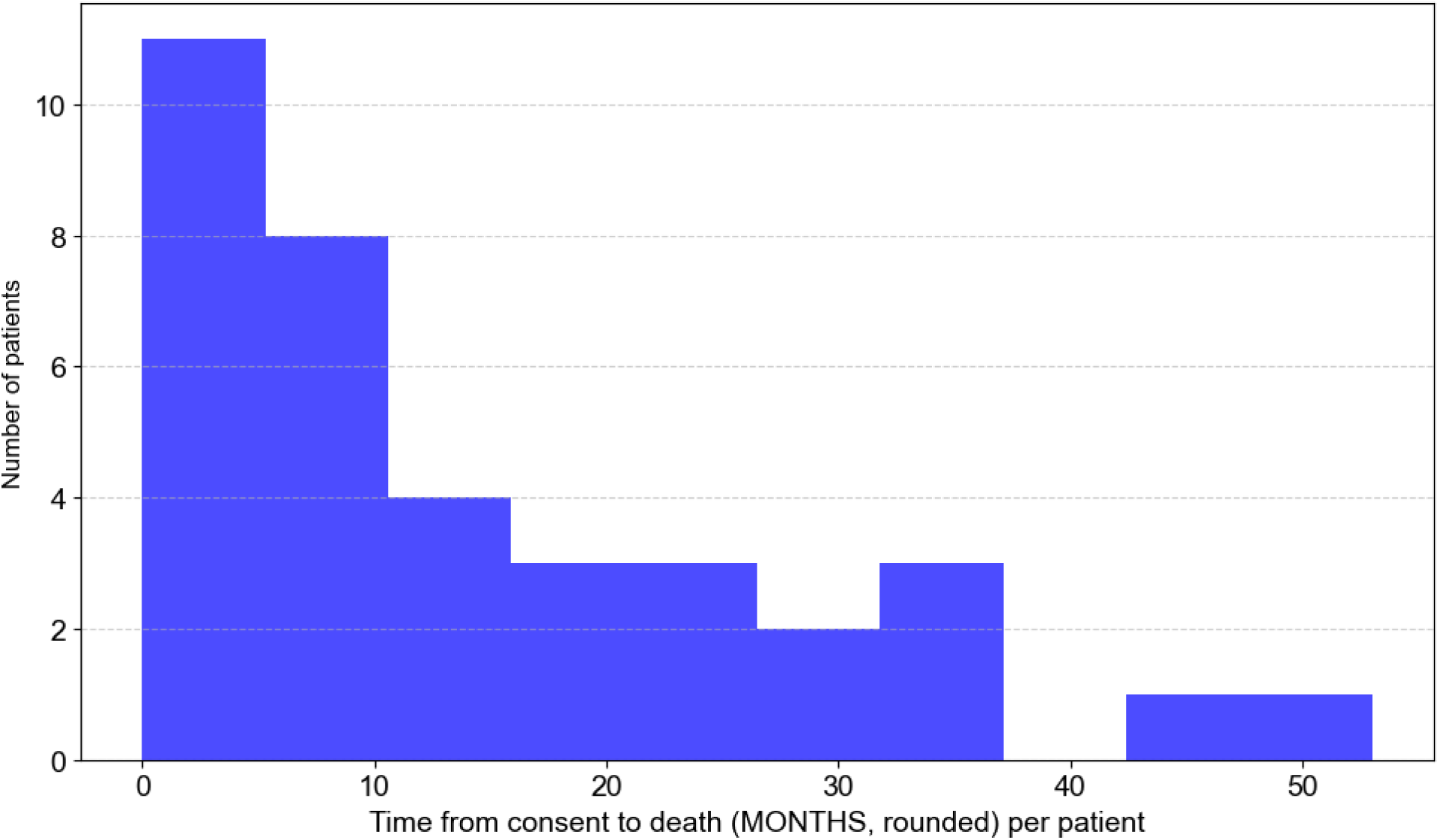
Histogram for time from consent to death per patient. Histogram summarizing our time from consent to death in months, rounded.

**Figure S2:**
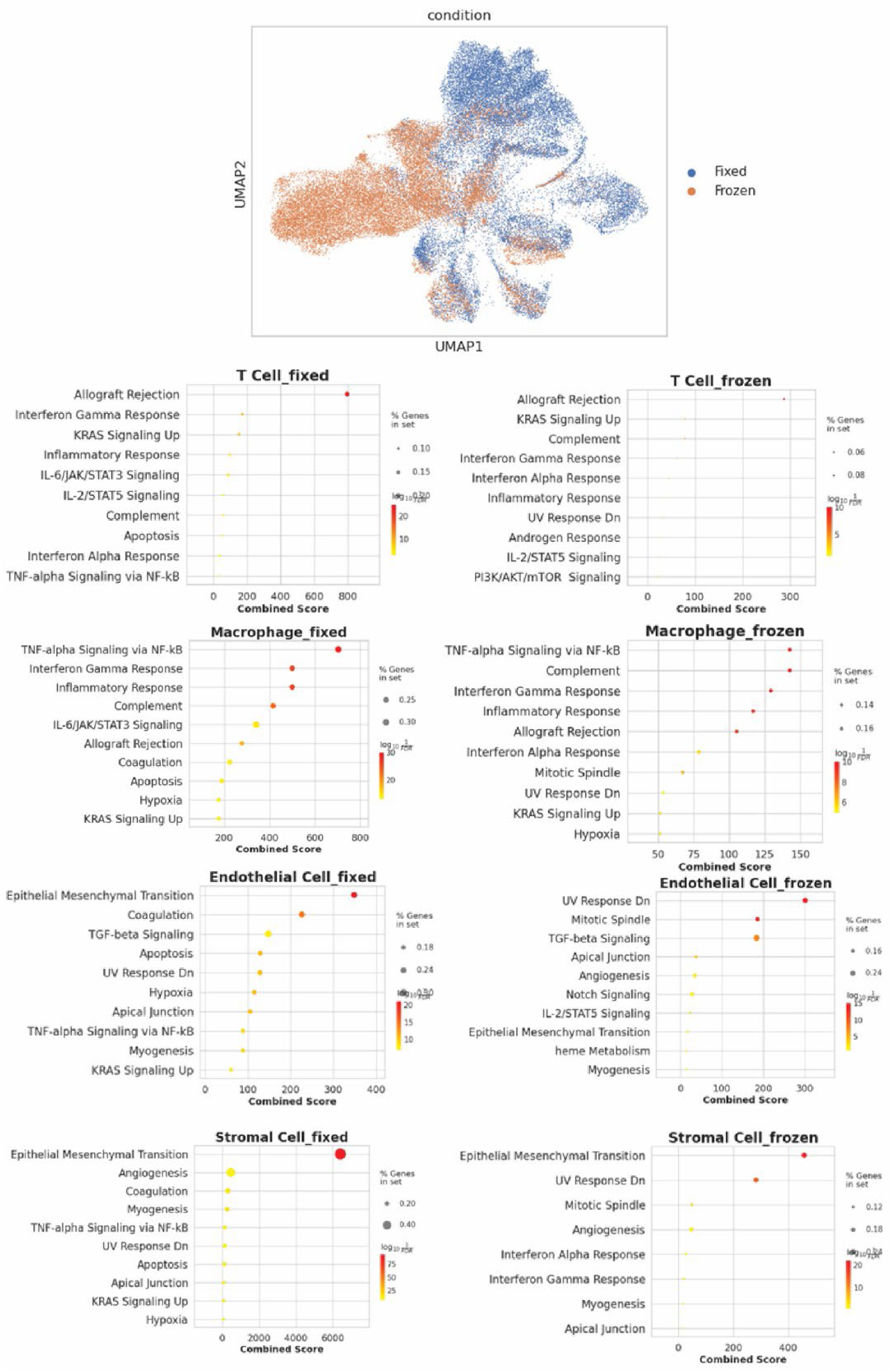
Pathway scores comparison between fixed and frozen tissue. Downstream single cell analysis showing that fixed sequencing has better gene set percentage overlap and higher scores on average.

**Figure S3:**
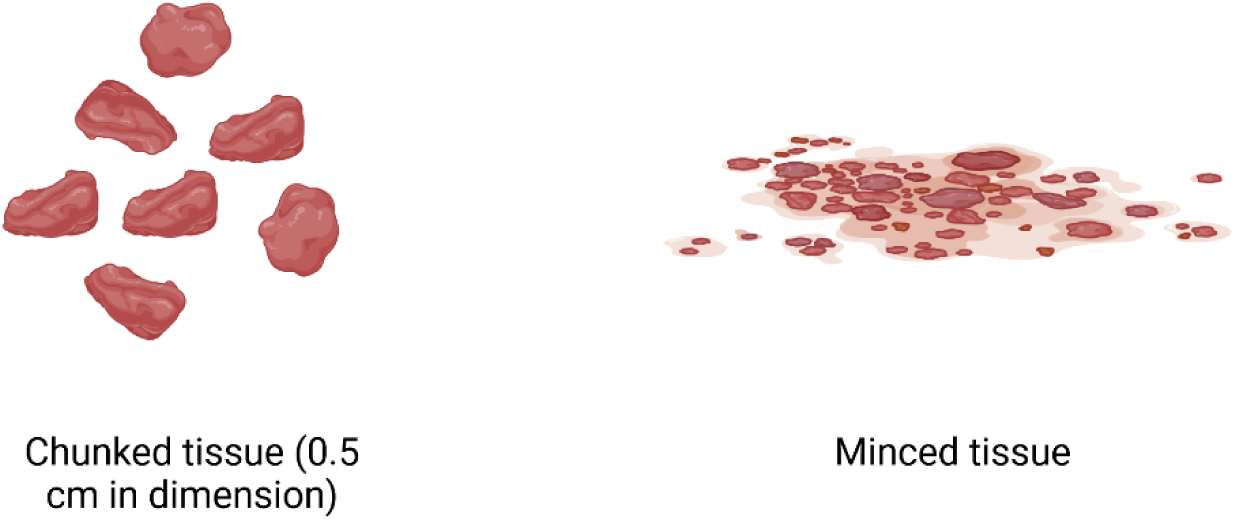
Illustration of difference between chunked and minced tissue. Illustration of the difference between chunked and minced tissue. Chunk tissues are preserved in 0.5cm – 1cm chunks. Minced tissues are chopped finely with a single-edged razor blade from an adjacent site.

## References

1. Breast Cancer Risk in American Women - NCI. December 17, 2020. Accessed August 2, 2024. https://www.cancer.gov/types/breast/risk-fact-sheet

2. Survival Rates for Breast Cancer. Accessed February 1, 2024. https://www.cancer.org/cancer/types/breast-cancer/understanding-a-breast-cancer-diagnosis/breast-cancer-survival-rates.html

3. Najjar MK, Manore SG, Regua AT, Lo HW. Antibody-Drug Conjugates for the Treatment of HER2-Positive Breast Cancer. Genes. 2022;13(11):2065. doi:10.3390/genes13112065

4. A Perspective on Cancer Cell Metastasis | Science. Accessed August 5, 2023. https://www.science.org/doi/10.1126/science.1203543

5. Dillekås H, Rogers MS, Straume O. Are 90% of deaths from cancer caused by metastases? Cancer Med. 2019;8(12):5574–5576. doi:10.1002/cam4.2474

6. Ren Q, Khoo WH, Corr AP, Phan TG, Croucher PI, Stewart SA. Gene expression predicts dormant metastatic breast cancer cell phenotype. Breast Cancer Res BCR. 2022;24:10. doi:10.1186/s13058-022-01503-5

7. Garcia-Recio S, Hinoue T, Wheeler GL, et al. Multiomics in primary and metastatic breast tumors from the AURORA US network finds microenvironment and epigenetic drivers of metastasis. Nat Cancer. 2023;4(1):128–147. doi:10.1038/s43018-022-00491-x

8. Duregon E, Schneider J, DeMarzo AM, Hooper JE. Rapid research autopsy is a stealthy but growing contributor to cancer research. Cancer. 2019;125(17):2915–2919. doi:10.1002/cncr.32184

9. Hooper JE. Rapid Autopsy Programs and Research Support: The Pre– and Post–COVID-19 Environments. Ajsp. 2021;26(2):100–107. doi:10.1097/PCR.0000000000000435

10. Rosenzweig M, Miller LA, Lee AV, et al. The Development and Implementation of an Autopsy/ Tissue Donation for Breast Cancer Research. New Bioeth Multidiscip J Biotechnol Body. 2021;27(4):349–361. doi:10.1080/20502877.2021.1993608

11. Lindell KO, Erlen JA, Kaminski N. Lessons from Our Patients: Development of a Warm Autopsy Program. PLOS Med. 2006;3(7):e234. doi:10.1371/journal.pmed.0030234

12. Desmedt C, Carey LA. Global post-mortem tissue donation programmes to accelerate cancer research. Nat Rev Cancer. 2024;24(5):289–290. doi:10.1038/s41568-024-00683-w

13. Achkar T, Wilson J, Simon J, Rosenzweig M, Puhalla S. Metastatic breast cancer patients: attitudes toward tissue donation for rapid autopsy. Breast Cancer Res Treat. 2016;155(1):159–164. doi:10.1007/s10549-015-3664-0

14. Robinson DR, Wu YM, Lonigro RJ, et al. Integrative Clinical Genomics of Metastatic Cancer. Nature. 2017;548(7667):297–303. doi:10.1038/nature23306

15. Aftimos P, Oliveira M, Irrthum A, et al. Genomic and Transcriptomic Analyses of Breast Cancer Primaries and Matched Metastases in AURORA, the Breast International Group (BIG) Molecular Screening Initiative. Cancer Discov. 2021;11(11):2796–2811. doi:10.1158/2159-8290.CD-20-1647

16. Boire A, Burke K, Cox TR, et al. Why do patients with cancer die? Nat Rev Cancer. 2024;24(8):578–589. doi:10.1038/s41568-024-00708-4

17. Kirwan CC, Descamps T, Castle J. Circulating tumour cells and hypercoagulability: a lethal relationship in metastatic breast cancer. Clin Transl Oncol. 2020;22(6):870–877. doi:10.1007/s12094-019-02197-6

18. Pereslucha AM, Wenger DM, Morris MF, Aydi ZB. Invasive Lobular Carcinoma: A Review of Imaging Modalities with Special Focus on Pathology Concordance. Healthcare. 2023;11(5):746. doi:10.3390/healthcare11050746

19. Oesterreich S, Pate L, Lee AV, et al. International survey on invasive lobular breast cancer identifies priority research questions. Npj Breast Cancer. 2024;10(1):1–7. doi:10.1038/s41523-024-00661-3

20. Geukens T, De Schepper M, Van Den Bogaert W, et al. Rapid autopsies to enhance metastatic research: the UPTIDER post-mortem tissue donation program. Npj Breast Cancer. 2024;10(1):1–14. doi:10.1038/s41523-024-00637-3

21. Single Cell Gene Expression Flex. 10x Genomics. Accessed February 8, 2024. https://www.10xgenomics.com/products/single-cell-gene-expression-flex

22. Guibert EE, Petrenko AY, Balaban CL, Somov AY, Rodriguez JV, Fuller BJ. Organ Preservation: Current Concepts and New Strategies for the Next Decade. Transfus Med Hemotherapy. 2011;38(2):125–142. doi:10.1159/000327033

23. Petrosyan V, Dobrolecki LE, LaPlante EL, et al. Immunologically “cold” triple negative breast cancers engraft at a higher rate in patient derived xenografts. NPJ Breast Cancer. 2022;8:104. doi:10.1038/s41523-022-00476-0

24. Hartmaier RJ, Trabucco SE, Priedigkeit N, et al. Recurrent hyperactive ESR1 fusion proteins in endocrine therapy-resistant breast cancer. Ann Oncol. 2018;29(4):872–880. doi:10.1093/annonc/mdy025

25. Lei JT, Shao J, Zhang J, et al. Functional Annotation of *ESR1* Gene Fusions in Estrogen Receptor-Positive Breast Cancer. Cell Rep. 2018;24(6):1434–1444.e7. doi:10.1016/j.celrep.2018.07.009

26. Yates ME, Waltermire H, Mori K, et al. ESR1 Fusions Invoke Breast Cancer Subtype-Dependent Enrichment of Ligand-Independent Oncogenic Signatures and Phenotypes. Endocrinology. 2024;165(10):bqae111. doi:10.1210/endocr/bqae111

27. Wang X, Zhou Y, Wu Z, et al. Single-cell transcriptomics reveals the role of antigen presentation in liver metastatic breast cancer. iScience. 2024;27(2). doi:10.1016/j.isci.2024.108896

28. Vallejo AF, Harvey K, Wang T, et al. snPATHO-seq: unlocking the FFPE archives for single nucleus RNA profiling. Published online December 6, 2022:2022.08.23.505054. doi:10.1101/2022.08.23.505054

