## Supplemental 1 - Copy of our SOP for "Hope for Others: Research Results from the University of Pittsburgh Rapid Autopsy Program for Breast Cancer"

Lee Oesterreich Lab

### Rapid Autopsy Lab SOP

### Standard Operating Procedure for the processing of patient specimens (solid tissue)

#### Morgue Protocol:

##### Supplies Needed in the morgue:

- 2 Ice chest/coolers filled with ice.
- 50ml tubes
- Marker pen
- Paper for listing samples
- 5 bottles of pre-chilled 500ml DMEM sample collection media
- 8 large 4 by 4 inch bitran bags for bone
- 8 large Ziploc dual sealed bags for organs.
- Bone marrow aspiration tool
- Liquid nitrogen dewar (if want to snap freeze in morgue)
- Optimal: two lab members and two separate vehicles.

| Organ | Number of Samples | Location |
|---|---|---|
| Brain | 3 | Frontal Lobe, Cerebellum, leptomeninges. |
| Lung | 5 | One from each lobe of the lung |
| Liver | 6 | Right tip, left tip, center, from the front and back of the organ |
| Primary Breast | 1 | Proximal to site of original primary |
| Contralateral breast | 1 | Mirrored to the site of original primary on the other side |

| | | |
|---|---|---|
| PE/Ascites | 2 | One from the pleural cavity, one from the abdominal cavity |
| Spleen | 1 | The whole spleen |
| Spine | 6 | Two cervical segment, two thoracic, two lumbar. |
| Any other metastatic sites | N/A | For ILC cases, we will also collect ovaries, omentum, peritoneum, uterus, and bladder (prioritize dome). |

Table 1: Organs collected and specific sites for tissue extraction.

##### *Morgue Processing*

Don PPE from the morgue, the Pathology Assistant, and the Resident can show you where to get it and how to put it on.

The autopsy then proceeds in the following order (conducted by the resident and assistant). For all of the following samples, record the number, site, and label on the autopsy tally sheet and autopsy map as you go. Request that all major organs have only peripheral small cuts removed for pathology department collections, and not bread-sliced, which is their usual practice. Put all samples on ice as soon as possible. Cover all samples with pre-chilled DMEM and bring back to the lab.

#### Lab Protocol:

##### Supplies Needed in the lab:

**Need 3-4 hoods depending on number of people available**

- Freezing media prepared (90% FBS, 10% DMSO)
- Cryovials
- Sterile surgical forceps
- Retractable scalpel
- Steak knives (if needed for huge liver samples)
- Bitran bags (small)
- 10% Neutral Buffered Formalin
- FFPE cassettes
- Pasteur pipettes
- Bleach
- Cavicide

- PPE-disposable gown, apron, gloves, and mask
- Oesteosoft
- 10 cm petri dish
- Possibly: Bitran bags, vacuum sealable for large tissue (looking into this)

#### Lab Processing

**<u>**Important- Please observe all appropriate BSL2+ procedures as human tumor samples are of unknown infectivity. Majority of these supplies can be found in the TC room cabinets underneath the centrifuge and sink, or on the RA shelf in the lab**</u>**

- Don disposable gown, apron, gloves, mask, and safety glasses.
- Double glove, so that you can easily dispose of the top pair and minimize exposure to potential pathogens if blood or other tissue contaminates top layer
- Avoid any glass- use pasteur pipettes when aspirating liquids
- Prepare beaker with 20% bleach to disinfect pipettes and all materials that come in contact with patient samples. Allow for disinfection time of 10 minutes.
- Clean TC hood and pipette gun, surgical tools with Cavicide, then clean a second time with 70% EtOH
- All associated trash should go into small red biohazard bags and then placed inside the regular red biohazard bins in the TC room (ie double bag everything)
- When handling specimen in the culture hood, use labelled petri dishes to contain the sample as you divide it, noting the overall size of the specimen

**NOTE: If multiple lesions are observed in a specific site and can be easily dissected from one another, these should be treated as individual specimens and clearly labelled in a way to easily track them.**

**Keep note of the time specimen was removed from patient, received by lab and when processing completed. As specimens are processed complete the 'Rapid Autopsy Tally sheet'.**

| Metadata | | Rapid Autopsy Number | Time of Death: | Autopsy Lab Members: | Autopsy Time: | Lab Processing Members: | Lab Processing Time: | Notes | This is post the SOP review in October 2023, by Alexander Chang | | | |
|---|---|---|---|---|---|---|---|---|---|---|---|---|
| DATA | | | | | | | | | | | | |
| Sample | Organ | Site | Gross dx (Tumor/ Normal) | Flash frozen (Lee/ Oesterreich) | FFPE | Flash frozen (PBC) | Cryo (minced) | Champions | Celcuity | In media for Daniel | Initials | Microscopic Tumor/Normal |
| 1 | | | | | | | | | | | | |
| 2 | | | | | | | | | | | | |
| 3 | | | | | | | | | | | | |
| 4 | | | | | | | | | | | | |
| 5 | | | | | | | | | | | | |
| 6 | | | | | | | | | | | | |
| 7 | | | | | | | | | | | | |
| 8 | | | | | | | | | | | | |
| 9 | | | | | | | | | | | | |
| 10 | | | | | | | | | | | | |
| 11 | | | | | | | | | | | | |
| 12 | | | | | | | | | | | | |
| 13 | | | | | | | | | | | | |
| 14 | | | | | | | | | | | | |
| 15 | | | | | | | | | | | | |

***<u>Prior to beginning any processing, observe a moment of silence out of respect for the patient and their donation.</u>***

*Organ Processing to extract tissues:*

For the large organs collected, we will collect and excise tissue samples from the locations identified in Table 1. Record on Autopsy Map accordingly, each extract tissue should go back on ice as soon as possible and be processed regularly.

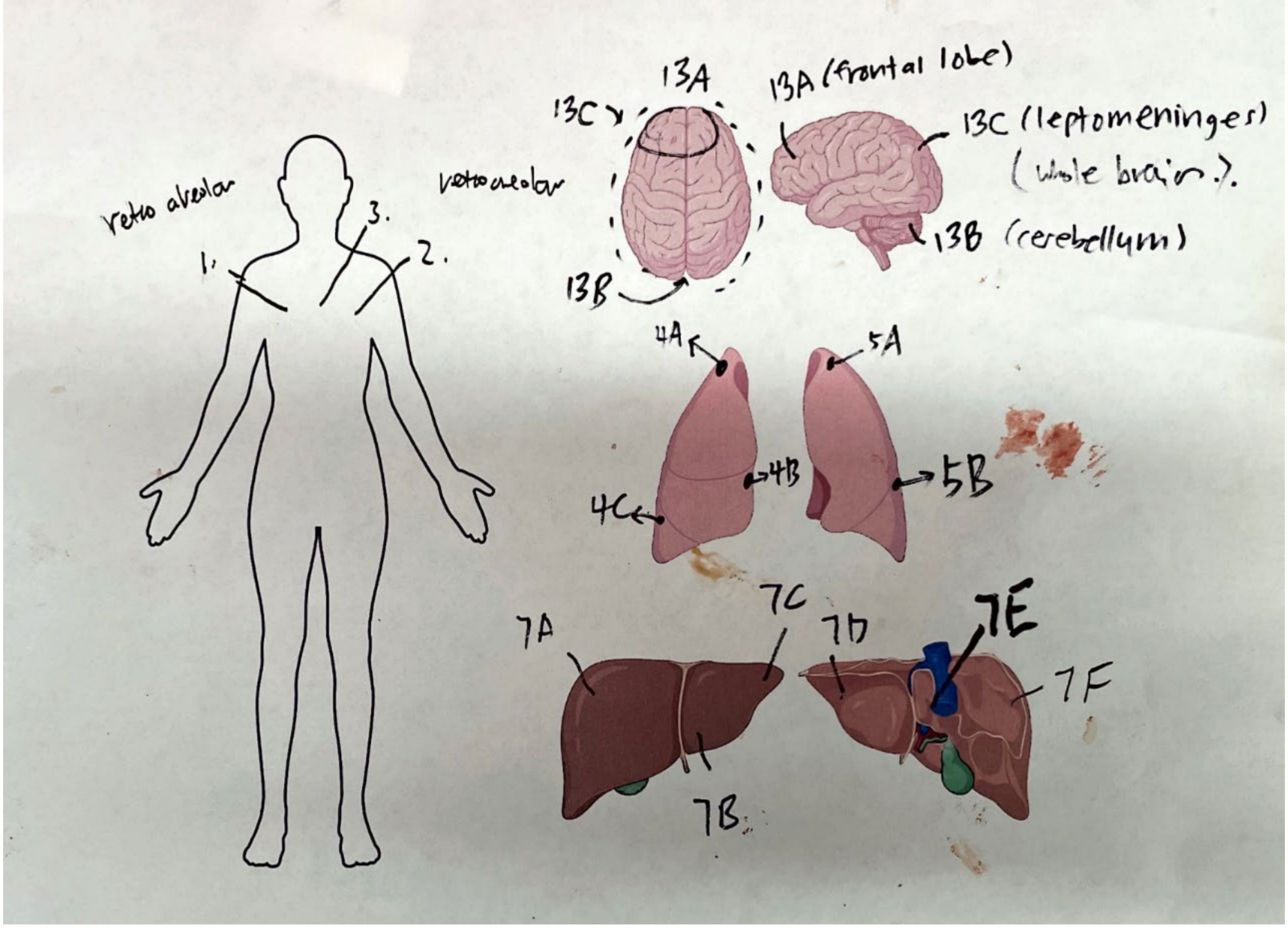


Figure 1 – Example Autopsy map.

*Tissue Processing:*

**Procedure listed in order of priority for fresh tumor tissue [to be confirmed with group ahead of each specific case] – <u>For each tissue (with the exception of bone) the goal is to collect (4 snap frozen bitran bags, 3 minced cryovials, 2 FFPE casettes)</u> :**

1. Snap freeze tumor and normal - Four samples. Priority is at least one for the lab (in hand labeled bitran bag). The second goes to PBC for banking and should be put in a bag with a barcode.
2. Formalin fix (FFPE) – two blocks for Lee-Oesterreich lab, more only if there is a lot of tissue.
3. For abundant sites, attempt organoid culture whenever possible.
4. If appropriate, tissue should be sent to Champions Oncology for PDX development (Lori will inform us of this is needed. If it is, we send the most abundant site(s).

5. Cryo preserve minced viable tissue (1ml FBS/10% DMSO and slow freeze) ~ 3 vials per site.

If excess tissue remains (after following the steps outlined), snap freeze an additional piece(s) and create extra FFPE blocks. Multiple pieces of the same tumor can go into bitran bags.

**1. Tissue priority #1: Snap frozen tissue**

a. Take a piece of tissue (size dependent on what is available) and place into a Bitran bag that has been labeled with the appropriate information pertaining to the sample (bar code for PBC, secureline marker for lab). Squeeze air out and seal bag.

b. Hold bag in liquid nitrogen for approximately one minute, submerging the sample but not the bag seal. Transfer to dry ice until storage. (note, if you submerge the bag in LN2 for more than a few minutes, when you remove the bag it will fill with gas and explode).

c. Store in –80 until future use.

Note: if tissue is a good size take frozen samples first for the lab (handwritten bitran bag, SecureLine marker) and then one for PBC (barcode bag)

**2. Tissue priority #2: Formalin fixation**

a. Take piece of tissue and place into a tissue cassette that has been labeled with the appropriate information pertaining to the sample i**n #2 pencil.** If the sample is very small, blue sponges are available to hold the sample inside the cassette. Note: only use pencil for labeling.

b. Place cassette with tissue in container with Formalin, 10% neutral buffer (Sigma) and leave O/N at 4°C.

c. Next day, switch to 70% EtOH and leave O/N at 4°C before taking to relevant histology team for paraffin embedding (research histology in Shadyside, $5 per sample for embedding).

**3. Tissue priority #3: Organoid development (for high priority cases)**

Follow the Lee Oesterreich PDO protocol to start organoid cultures from as many sites as feasible.

**4. Tissue priority #4: To Champions oncology, if appropriate**

Follow Champions oncology requirements for shipping small tumor specimen in media. Lori will let the lab know if samples should be sent to Champions. If they are to be sent, we have tubes of media from Champions in which we should put 2-3 small tumor fragments (approx. 2x5mm). Lori will instruct on shipping the samples.

**5. Tissue priority #5: cryopreservation of viable minced tissue for later organoid preparation**

a. Prepare freezing media (90% FBS, 10% DMSO)

b. Mince tissue into tiny chunks, as small as possible with scalpel blades (approx. 1x 1mm) in cryovial and pipette 1 ml of freezing media into tube, the equivalent material to multiple small chunks per vial. Invert tube a few times to mix.

c. Ensure tubes are labelled appropriately

d. Place tubes in ‘Mr frosty’ or Styrofoam box to allow for slow cooling, and place in -80c freezer. The samples should go to the freezer within 5 minutes (not sit on counter until its full)

e. After 24 hours, store long-term in LN2 until future use, completing a box map of the contents for lab documentation

**6. Bone samples:**

1. Take one half of the pre-bisected sample and put in small bitran bag for flash freezing, take the other and put it in FFPE cassette and proceed with below steps.
2. Fix in Formalin for 2 days (4C).
3. Switch to Osteosoft and incubate with shaking in 4C.
4. Most bones might take 7 days – maybe 10 days for spine.
5. Switch to 70% ethanol.
6. Send for Paraffin embedding.

**Documentation and storage at time of autopsy processing:**

- As samples are processed, the sample ‘tally sheet’ should be completed.
- Frozen samples should be separated so that one box is prepared for PBC with one snap frozen sample of each site (in barcoded bags). Other snap frozen samples are for lab. Ensure all boxes are labeled on the top and side with the TP number of the autopsy and what sample type is inside (ie, Flash frozen tissue).

**Clean up:**

- Clean TC hood and pipette gun, surgical tools with Cavicide, then clean a second time with 70% EtOH
- All associated trash should go into small red biohazard bags and then placed inside the regular red biohazard bins in the TC room (so waste is double bagged).
- All surgical tools should be soaked in a 20% bleach solution overnight.

**Documentation and storage the day after the autopsy:**

- Formalin fixed samples should be transferred to 70% ethanol. After 24 hours in ethanol, samples should be sent to PBC/research histology (Shadyside) for parrafin embedding (wax blocks). Processed samples are inventoried and stored in Assembly Room 1615 (Jenny's old office).
- Frozen samples for PBC should be provided to PBC, alongside the following form which should be completed:
- Frozen samples for the lab should be stored in an appropriate location and updated in the lab -80 or -150 inventory (tumor –80 freezer, and special rack)
- Cyrovials in the Mr Frosty or styrofoam containers should be transferred to a freezer box for LN2 storage. Make sure the box is labeled with the TP number and 'rapid autopsy'. A box map for the LN2 viable cryo samples must be made and added to both the patient folder on onedrive, and to the lab LN2 inventory spreadsheet.
- The sample log which was completed during autopsy processing should be typed up into the form and added to the patient folder on one drive. Any other notable things from the autopsy or processing should also be recorded and saved to the OneDrive folder.
- Surgical tools should be rinsed in water, cleaned with ethanol, and sent for sterilization. Once sterilized they should be stored for the next autopsy.
